# CARD9 in Neutrophils Protects from Colitis and Controls Mitochondrial Metabolism and Cell Survival

**DOI:** 10.1101/2022.01.14.476327

**Authors:** Camille Danne, Chloé Michaudel, Jurate Skerniskyte, Julien Planchais, Aurélie Magniez, Allison Agus, Marie-Laure Michel, Bruno Lamas, Gregory Da-Costa, Madeleine Spatz, Cyriane Oeuvray, Chloé Galbert, Maxime Poirier, Yazhou Wang, Alexia Lapiere, Nathalie Rolhion, Tatiana Ledent, Cédric Pionneau, Solenne Chardonnet, Floriant Bellvert, Edern Cahoreau, Amandine Rocher, Rafael Jose Argüello, Carole Peyssonnaux, Sabine Louis, Mathias L. Richard, Philippe Langella, Jamel El-Benna, Benoit Marteyn, Harry Sokol

## Abstract

**Objectives:** Inflammatory bowel disease (IBD) results from a combination of genetic predisposition, dysbiosis of the gut microbiota and environmental factors, leading to alterations in the gastrointestinal immune response and chronic inflammation. Caspase recruitment domain 9 (*Card9*), one of the IBD susceptibility genes, has been shown to protect against intestinal inflammation and fungal infection. However, the cell types and mechanisms involved in the CARD9 protective role against inflammation remain unknown.

**Design:** We used dextran sulfate sodium (DSS)-induced and adoptive transfer colitis models in total and conditional CARD9 knock-out mice to uncover which cell types play a role in the CARD9 protective phenotype. The impact of *Card9* deletion on neutrophil function was assessed by an *in vivo* model of fungal infection and various functional assays, including endpoint dilution assay, apoptosis assay by flow cytometry, proteomics and real time bioenergetic profile analysis (Seahorse).

**Results:** Lymphocytes are not intrinsically involved in the CARD9 protective role against colitis. CARD9 expression in neutrophils, but not in epithelial or CD11c+ cells, protects against DSS-induced colitis. In the absence of CARD9, mitochondrial dysfunction in neutrophils leads to their premature death through apoptosis, especially in oxidative environment. The decrease of fonctional neutrophils in tissues could explain the impaired containment of fungi and increased susceptibility to intestinal inflammation.

**Conclusion:** These results provide new insight into the role of CARD9 in neutrophil mitochondrial function and its involvement in intestinal inflammation, paving the way for new therapeutic strategies targeting neutrophils.

**Summary box:**

1. **What is already known about this subject?**
  - Inflammatory bowel disease (IBD) results from genetic predisposition, microbiota dysbiosis and environmental factors, but the alterations of the immune response leading to chronic intestinal inflammation are still not fully understood.
  - Caspase recruitment domain 9 (*Card9*), one of the IBD susceptibility genes, has been shown to protect against intestinal inflammation and fungal infection.
  - However, the cell types and cellular mechanisms involved in the CARD9 protective role against inflammation remain unknown.
2. **What are the new findings?**

- CARD9 expression in neutrophils, but not in lymphocytes, epithelial cells or CD11c+ cells, protects against DSS-induced colitis.
- In the absence of CARD9, mitochondrial dysfunction in neutrophils leads to their premature death through apoptosis, especially in oxidative environment.
- The decrease of fonctional neutrophils in tissues could explain the impaired containment of fungi and increased susceptibility to intestinal inflammation.
3. **How might it impact on clinical practice in the foreseeable future?**
  - These results provide new insight into the role of CARD9 in neutrophil mitochondrial function and its involvement in intestinal inflammation.
  - Understanding the role of neutrophils in chronic inflammation could lead to innovative therapeutic strategies targeting these key immune cells for various complex diseases.

## INTRODUCTION

Inflammatory bowel disease (IBD) results from a combination of genetic predisposition, dysbiosis of the gut microbiota and environmental factors, leading to alterations in the gastrointestinal immune response and chronic inflammation^1,2^. Especially, the innate compartment of the immune system has been involved in IBD development, with a role for dendritic cells, macrophages and neutrophils^3–6^. Neutrophils, one of the most abundant and important mediators of innate immunity, are professional phagocytes that mount the acute inflammatory response and act as the first line of defense against invading pathogens^7,8^. An impaired neutrophil function may result in limited pathogen clearance and fuel a chronic inflammatory response with excessive lymphocyte activation. Patients with congenital disorders in neutrophil function such as chronic granulomatous disease (CGD) often develop IBD-like phenotypes^9–12^. Moreover, functional defects have been observed in neutrophils from IBD patients, including impaired chemotaxis, migration, phagocytosis or ROS production^4,5^.

CARD9, one of the numerous IBD susceptibility genes, encodes an adaptor protein that integrates signals downstream of pattern recognition receptors ^13–18^. Especially, CARD9 is involved in the host defense against fungi via C-type lectin sensing^19,20^. CARD9 polymorphisms in humans are associated with multiple susceptibilities including IBD^21^, whereas loss-of-function mutations are associated with invasive fungal infections caused by species such as *Candida albicans*^21–24^. CARD9 was shown to mediate its protective functions, at least in part, through the induction of adaptive Th17 cell responses^22,23,25^. *Card9*^-/-^ mice are more susceptible to colitis due to impaired IL-22 production and have an increased load of gut-resident fungi^25^. Indeed, CARD9 affects the composition and function of the gut microbiota, altering the production of anti-inflammatory microbial metabolites^26,27^. However, the cell types involved in the CARD9 protective role against intestinal inflammation remain unknown.

In this work, we show that CARD9 expression in neutrophils, but not in epithelial or CD11c+ cells such as dendritic cells, protects against dextran sulfate sodium (DSS)-induced colitis in mice. The absence of CARD9 impacts neutrophil capacity to contain fungal dissemination, notably by impairing neutrophil mitochondrial function and survival. Indeed, *Card9* deletion induces a basal overactivation of mitochondria, increasing mitochondrial dysfunction and apoptosis in neutrophils. These results provide new insight into the role of CARD9 in neutrophil mitochondrial function and its consequences in intestinal inflammation.

## RESULTS

### Lymphocytes have no intrinsic role in the *Card9^-/-^* susceptibility to colitis

CARD9 was previously reported to be mainly expressed in myeloid cells, especially dendritic cells, macrophages and neutrophils^14,28^. Using qRT-PCR analyses in various C57BL/6 mouse organs, we confirmed that *Card9* is mainly expressed in immune organs such as bone marrow, spleen and distal small intestine (ileum), but is low at baseline in proximal and mid small intestine, caecum, colon, stomach and liver, and not detectable in *Card9*^-/-^ tissues (Fig. S1A). Consistently, western-blot analysis showed the expression of CARD9 protein in bone marrow, spleen and distal small intestine of WT mice (Fig. S1B). To dissect *Card9* expression at the cellular level, we sorted immune cell populations from spleen and bone marrow of WT mice. *Card9* is highly expressed in neutrophils (Ly6G^+^CD11b^+^ cells), macrophages (CD11b^+^F4/80^+^ cells), CD11c^+^ cells, including dendritic cells, and monocytes (CD11b^hi^F4/80^+^ cells); but the expression is lower in innate or adaptive lymphocytes (CD3^+^TCRγδ^+^ lymphoid cells, CD3^+^CD4^+^ and CD3^+^CD8^+^ T cells, CD3^-^CD19^+^ B cells) (Fig. S1C). Thus, CARD9 likely plays a major role within the myeloid immune compartment.

Previous studies from our group and others started to investigate the role of *Card9* in murine models of experimental colitis, showing that *Card9* deletion increases colitis susceptibility^25,26,29^. In order to decipher the respective roles of lymphocytes and myeloid cells in the *Card9* susceptibility to intestinal injury and inflammation, we first induced colitis with DSS in *Rag2*^-/-^ and *Rag2*^-/-^x*Card9*^-/-^ mice that are deficient in functional T and B cells. Mice were euthanized after receiving 3% DSS in drinking water for 7d, as the severity limit was reached for the *Rag2*^-/-^x*Card9*^-/-^ group. Indeed, desease severity (defined by weight loss, DAI (Disease Activity Index) score, colon length, and histologic score) was strongly increased in *Rag2*^-/-^x*Card9*^-/-^ mice compared to *Rag2*^-/-^ mice (Fig. 1A-E). In an adoptive transfer model of colitis, in which *Rag2*^-/-^ mice lacking functional lymphocytes received T cells either from WT or *Card9*^-/-^ mice, no difference was observed on colitis severity (Fig. 1F), meaning that *Card9* expression in T cells does not impact colitis susceptibility. However, the transfer of WT T cells into *Rag2*^-/-^x*Card9*^-/-^ recipient mice did aggravate colitis compared to *Rag2*^-/-^ simple KO recipients, with a significantly stronger weight loss (Fig. 1F). These results demonstrate that *Card9* mediates its protective role against colitis through the innate immunity compartment, although its role in intestinal epithelial cells cannot be ruled out.

**Figure 1.**
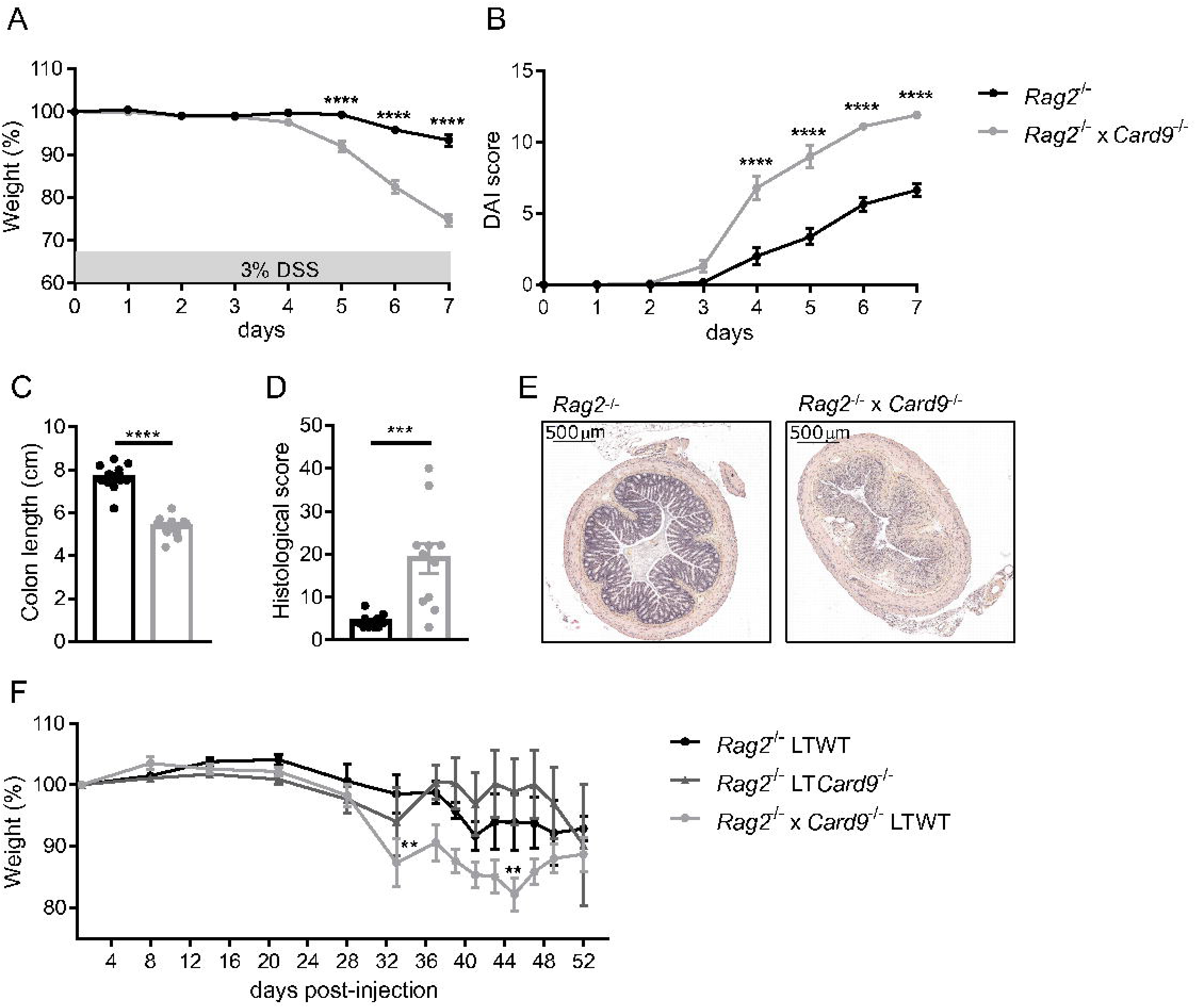
Lymphocytes have no intrinsic role in the *Card9*^-/-^ phenotype in colitis models. (A) Weight and (B) Disease Activity Index (DAI) score of DSS-exposed *Rag2*^-/-^ or *Rag2^-/-^ x Card9*^-/-^ mice. (C) Colon length and (D) histological score of colon sections at day 7. Data points represent individual mice. One representative experiment out of three. (E) Representative H&E-stained images of mid colon cross-sections from DSS-exposed *Rag2*^-/-^ (left) and *Rag2^-/-^ x Card9*^-/-^ (right) mice at day 7. Scale bars, 500 μm. (F) Weight of mice receiving naive T cells for adoptive transfer of colitis experiment. *Rag2*^-/-^ received either WT or *Card9*^-/-^ lymphocytes. *Rag2^-/-^ x Card9*^-/-^ mice received WT lymphocytes. Data are mean ± SEM of two independent experiments. *P<0.05, **P<0.01, ***P<0.001, ****P<0.0001 as determined by two-way analysis of variance (ANOVA) with Sidak’s post-test (A, B, F) and Mann-Whitney test (C, D). LT, lymphocytes T.

### *Card9* expression in neutrophils, but not in epithelial or CD11c^+^ cells, protects against colitis

Based on these findings, we generated conditional KO mice using the cre-lox technology, and obtained mouse strains defective for *Card9* either in epithelial cells only (Villincrex*Card9*lox line), CD11c-expressing cells only, including dendritic cells, macrophages and monocytes (CD11ccrex*Card9*lox line), or neutrophils only (Mrp8crex*Card9*lox line). To validate their phenotypes, we isolated epithelial cells from the colonic lamina propria of *Card9*^Villincre^ and *Card9*^Villinwt^ mice, or used MACS separation columns to isolate either CD11c^+^ or Ly6G^+^ cell fractions from spleen or bone marrow of *Card9*^CD11cwt^ and *Card9*^CD11ccre^ mice or *Card9*^Mrp8wt^ and *Card9*^Mrp8cre^ mice, respectively. We performed qRT-PCR on these cell fractions (Fig. S2A), and western-blot analyses on the Ly6G^+^ and Ly6G^low/-^ fractions of *Card9*WT, *Card9^-/-^, Card9*^Mrp8wt^ and *Card9*^Mrp8cre^ mice (Fig. S2B). Results confirmed that *Card9* deletion is restricted to the expected cell types (Fig. S2A-B). Purity of Ly6G^+^CD11b^+^ neutrophils isolated from the bone marrow of C57Bl/6 mice reached 95% by flow cytometry (Fig. S2C).

We then assessed the susceptibility of these newly generated mice strains in a model of intestinal inflammation. DSS was administered for 7 days, followed by additional 5 days in which DSS was discontinued. The deletion of *Card9* in epithelial or CD11c^+^ cells did not affect mouse susceptibility to colitis (Fig. 2A-B and S2D). However, the deletion of *Card9* in neutrophils aggravates colitis compared to WT littermate controls, with significantly increased weight loss from day 8 (Fig. 2C), DAI score from day 5 (Fig. 2C), and histological score (Fig. 2E-F), as well as decreased colon length (Fig. 2D). Thus, the expression of *Card9* in neutrophils plays a crucial role in the protection against intestinal inflammation. The expression of myeloperoxidase (MPO), an anti-microbial enzyme abundantly expressed in neutrophils, was increased in *Card9*^Mrp8cre^ compared to *Card9*^Mrp8wt^ colon tissue at day 12, indicating a more important presence or activation of neutrophils in the absence of *Card9* (Fig. 2G). Similarly, the expression of the inflammatory marker lipocalin (*Lcn2*) was increased in *Card9*^Mrp8cre^ colon tissue at day 12 (Fig. 2H).

**Figure 2.**
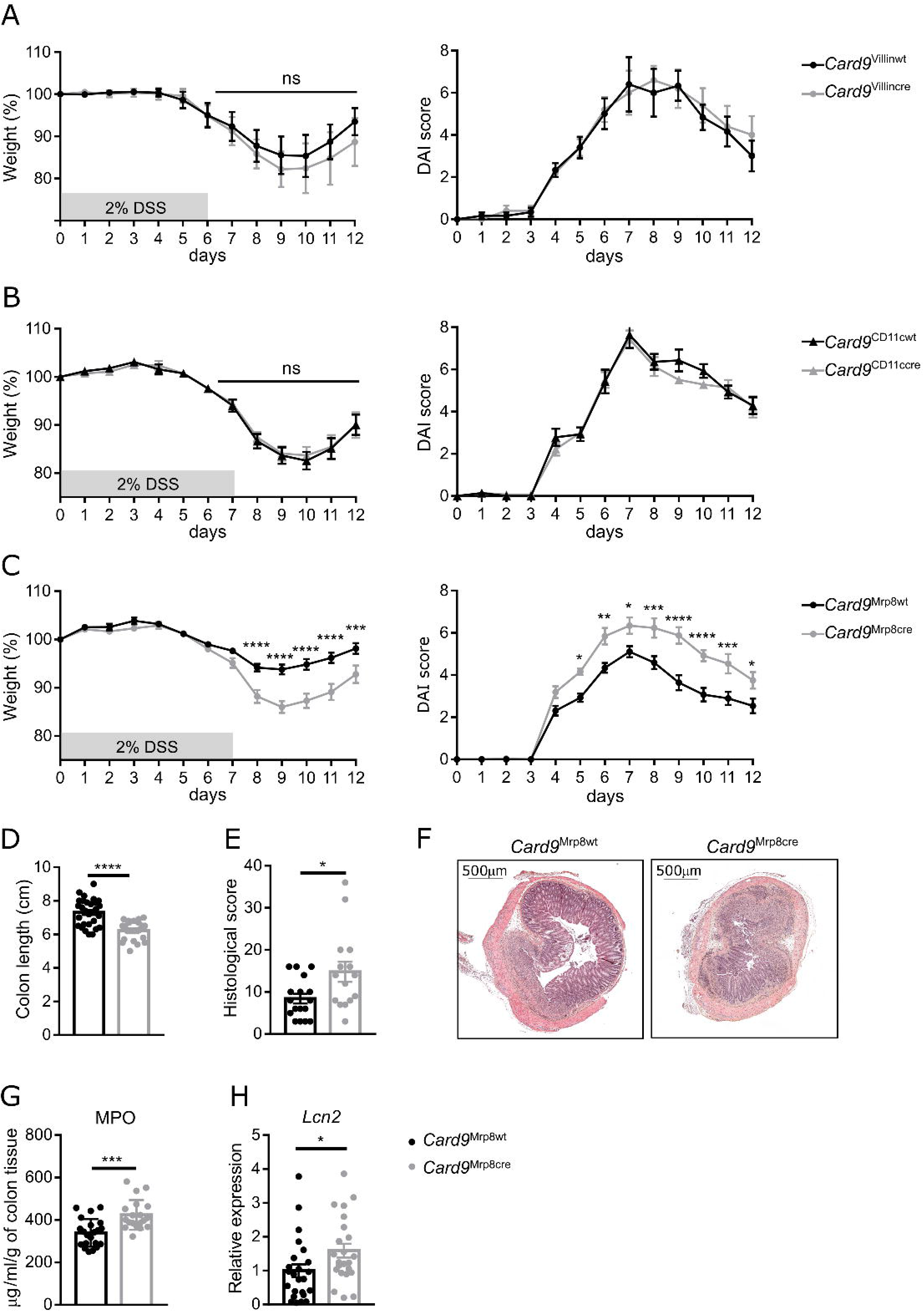
*Card9* expression in neutrophils, but not in epithelial or CD11c^+^ cells, protects against colitis. (A) Weight and DAI score of DSS-exposed *Card9*^Villinwt^ or *Card9*^Villincre^ mice. (B) Weight and DAI score of DSS-exposed *Card9*^cd11cwt^ or *Card9*^cd11ccre^ mice. (C) Weight and DAI score of DSS-exposed *Card9*^Mrp8wt^ or *Card9*^Mrp8cre^ mice. (D) Colon length, (E) histological score of colon sections and (F) representative H&E-stained images of mid colon cross-sections from DSS-exposed *Card9*^Mrp8wt^ (left) and *Card9*^Mrp8cre^ (right) mice at day 12. Scale bars, 500 μm. (G) Myeloperoxidase (MPO) concentration in total colon tissue at day 12. (H) Lipocalin (*Lcn2*) expression by qRT-PCR in total colon tissue at day 12, normalized to *Gapdh*. Data points represent individual mice. Data are mean ± SEM of three independent experiments. *P<0.05, **P<0.01, ***P<0.001, ****P<0.0001 as determined by two-way ANOVA with Sidak’s post-test (A-C) and Mann-Whitney test (D-H).

These results suggest that *Card9*-deficient neutrophils are efficiently recruited to the inflamed tissue but likely exhibit functional defects preventing them from adequately controlling microbial invaders and thus maintaining inflammation within the intestinal mucosa. We previously showed that the gut microbiota of total *Card9*^-/-^ KO lineage mice exhibit an altered production of AhR agonists compared to WT^26^. However, no difference was observed between *Card9*^Mrp8wt^ and *Card9*^Mrp8cre^ mice (Fig. S2E), suggesting that this aspect of the phenotype is not intrinsically related to the role of CARD9 in neutrophils.

### *Card9* deletion affects the number of activated neutrophils in the inflamed colon

To investigate neutrophil function in WT mice during DSS-induced inflammation, we analyzed RNA expression of the neutrophil-specific genes *Lcn2, Cxcr2* and *S100A8* in colon tissue at days 0, 4, 7, 9, 12 and 16 of DSS-induced colitis (Fig. 3A and S2F). The neutrophil recruitment was maximal at day 9, corresponding to the peak of clinical inflammation, and remained high up to day 16 (Fig. 3A). Histological sections of the distal colon confirmed these findings (Fig. 3B). The fact that inflammation was higher in *Card9*^Mrp8cre^ than in *Card9*^Mrp8wt^ mice (Fig. S2E), and that *Lcn2, Cxcr2* and *S100a8* expression at day 9 in colon tissue were similar in both genotypes (Fig. 3C) excludes the hypothesis of a defect of neutrophil recruitment in *Card9*^Mrp8cre^ mice. We then examined the immune cell populations recruited to the colon lamina propria (LP) of *Card9*^Mrp8cre^ versus *Card9*^Mrp8wt^ mice at day 9 of colitis. Surprisingly, although the total number of neutrophils was similar in the two genotypes, the percentage and count of mature and activated Ly6G^+^CD11b^+^ neutrophils was decreased in the colon LP of *Card9*^Mrp8cre^ (Fig. 3D-E). Moreover, the expression levels of both Ly6G and CD11b surface proteins, two major neutrophil maturationa and activation markers, were significantly reduced in the overall neutrophil population, as shown by decreased MFIs (for Mean Fluorescence Intensity) (Fig. 3F-G). These findings suggest a structural or functional defect in neutrophils deleted for *Card9*.

**Figure 3.**
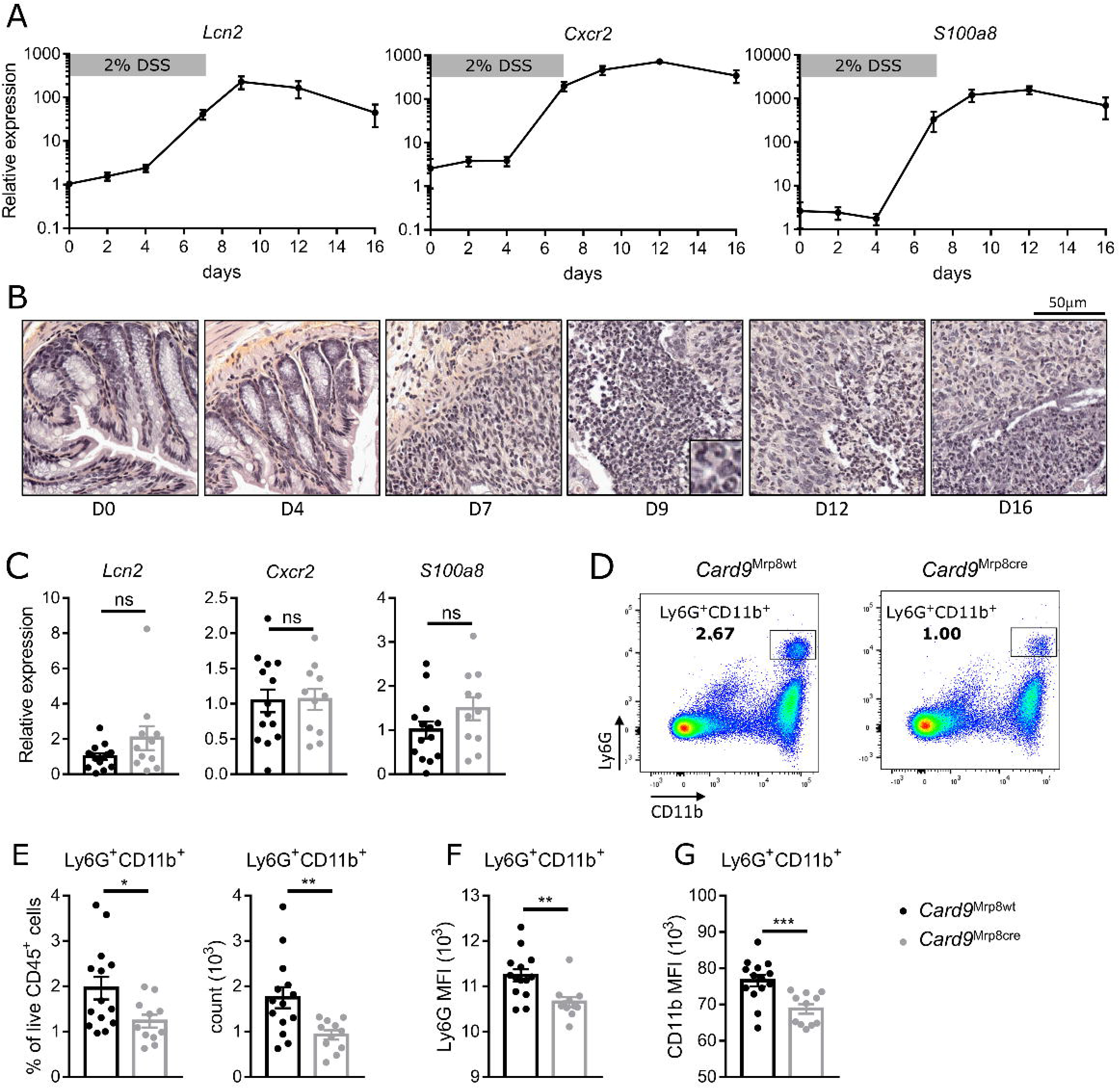
*Card9* deletion affects the number of neutrophils in the inflamed colon. (A) Relative expression of *Lcn2, Cxcr2* and *S1008a* in distal colon tissue of C57BL/6 WT mice during a DSS colitis model relative to *Gapdh*. (B) Representative H&E-stained images of mid colon cross-sections from DSS-exposed WT mice at day 0, 4, 7, 9, 12 and 16 after DSS exposure. Scale bars, 50 μm. (C) *Lcn2, Cxcr2* and *S100A8* expression in total colon tissue from *Card9*^Mrp8wt^ and *Card9*^Mrp8cre^ mice at day 9 of DSS colitis by qRT-PCR analyses, normalized to *Gapdh*. (D) Representative flow cytometry plots of Ly6G^+^CD11b^+^ cells (neutrophils) in the colon lamina propria (LP) of DSS-exposed *Card9*^Mrp8wt^ and *Card9*^Mrp8cre^ mice. (E) Percentage and count of Ly6G^+^CD11b^+^ cells in the LP of DSS-exposed *Card9*^Mrp8wt^ and *Card9*^Mrp8cre^ mice. (F) Ly6G and (G) CD11b expression (MFI for Mean Fluorescence Intensity) of Ly6G^+^CD11b^+^ neutrophils from *Card9*^Mrp8wt^ and *Card9*^Mrp8cre^ mice. Data points represent individual mice. Data are mean ± SEM of two independent experiments. *P<0.05, **P<0.01, ***P<0.001 as determined by Mann-Whitney tests.

### *C. albicans* killing capacities are impacted by *Card9* deletion in neutrophils

To investigate the defect caused by *Card9* deletion in *Card9*^Mrp8cre^ compared to *Card9*^Mrp8wt^ neutrophils, we developed several *in vitro* assays with Ly6G^+^ neutrophils purified from mouse bone marrow using MACS separation columns. Immunofluorescence, scanning electron microscopy (SEM) and transmission electron microscopy (TEM) analyses did not reveal noticeable structural differences between both genotypes (Fig. S3). On the functional side, we tested the neutrophils ability to kill microorganisms, especially fungi, as CARD9 plays a crucial role in host defense against fungal infection in both humans and mice^25,20^. Indeed, *C. albicans* killing capacities are strongly affected by *Card9* deletion in neutrophils (Fig. 4). An endpoint-dilution survival assay in 96-well plates revealed that twice as many *C. albicans* cfu (colony forming unit) do survive after 24h co-incubation with *Card9*^-/-^ or *Card9*^Mrp8cre^ neutrophils compared to Card9WT or *Card9*^Mrp8wt^ controls, respectively (Fig. 4A-B). A killing assay using the cfu counting method on agar plates confirmed that *Card9*^Mrp8cre^ neutrophils have impaired abilities to kill *C. albicans* compared to *Card9*^Mrp8wt^ (Fig. 4C). *Card9* deletion in neutrophils did not impact phagocytosis *per se*, as shown by flow cytometry experiments using fluorescein-isothiocyanate (FITC)-conjugated zymosan (from *Saccharomyces cerevisiae* cell wall), or cultures of live *C. albicans-GFP* or *E. coli-GFP* (Fig. S4A-B). Moreover, we did not observe a difference in the levels of Reactive Oxygen Species (ROS) production over time between neutrophils of the two genotypes in response to phorbol myristate acetate (PMA), a PKC-dependent neutrophil activator, or zymosan, a fungal stimulus (Fig. S4C-E). Even though *Card9* is involved in autophagy^30^, this cellular process did not seem to be affected by *Card9* deletion in neutrophils *in vitro*, as shown by the normality of p62 and LC3BII/I ratio on western-blots (Fig. S4F).

**Figure 4.**
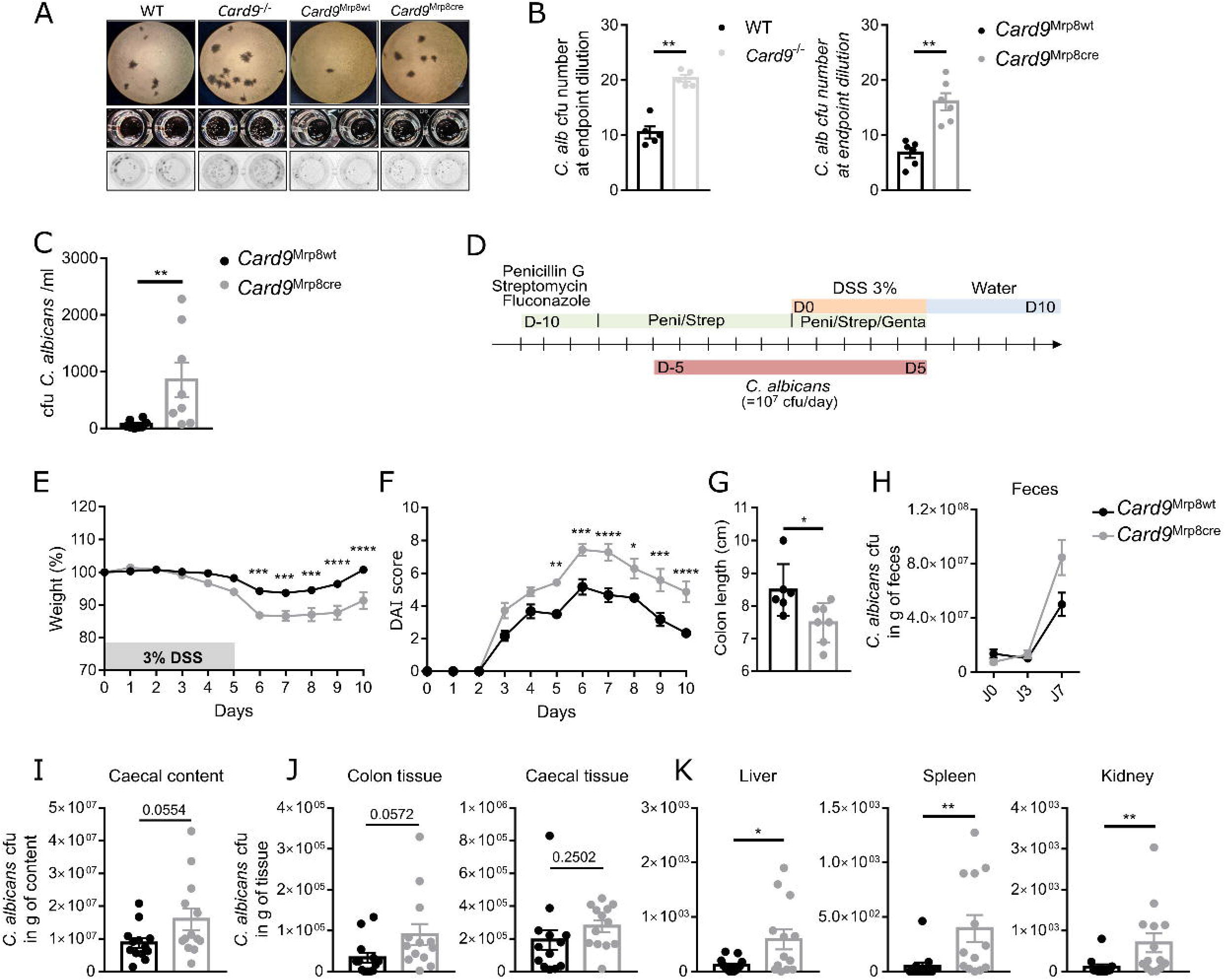
*C. albicans* killing capacity is impacted by *Card9* deletion in neutrophils. (A) Representative images of *C. albicans* cfu number at the endpoint dilution after infection of *Card9*WT, *Card9^-/-^, Card9*^Mrp8wt^ or *Card9*^Mrp8cre^ neutrophils for 24h, using a microscope (top) or directly showing *C. albicans* cfu in the 96-well plate, with a photography from above (middle picture) or a scan from below the plate (bottom picture). (B) *C. albicans* cfu after infection of neutrophils for 24h. Cfu counting was performed in 96 well plates using a microscope. Data points represent individual mice. Data are mean ± SEM of three independent experiments. (C) *C. albicans* cfu number after infection of *Card9*^Mrp8wt^ or *Card9*^Mrp8cre^ neutrophils for 3 or 24h. Cfu counting was performed after plating on YEPD agar plates. Data are mean ± SEM of two independent experiments. (D) Experimental design of antibiotic treatments and *C. albicans* inoculation in mice treated with 3% DSS. (E) Weight, (F) DAI and (G) colon length of DSS-exposed *Card9*^Mrp8wt^ and *Card9*^Mrp8cre^ mice after colonization with *C. albicans*. (H-K) Fungal burden in the feces (H), caecal content (I), caecal and colon tissue (J) and liver, spleen and kidney (K) of DSS-exposed *Card9*^Mrp8wt^ and *Card9*^Mrp8cre^ mice after colonization with *C. albicans*. Data are mean ± SEM of two independent experiments. *P<0.05, **P<0.01, ***P<0.001, ****P<0.0001 as determined by one-way ANOVA with Tukey’s post-test (A), two-way ANOVA with Sidak’s post-test (E, F) and Mann-Whitney test (C, G, I-K). Genta, gentamycin.

We thus followed the track of fungal killing to investigate whether the absence of *Card9* expression in neutrophils drives a general impairment of the immune system to control infection. During a DSS-induced colitis model, we detected a slight but non-significant increase in total fungi/bacteria DNA ratio at days 7 and 12 in the feces of *Card9*^Mrp8cre^ compared to *Card9*^Mrp8wt^ mice (Fig. S4G). These data suggest a reduced ability of *Card9*^Mrp8cre^ mice to contain fungi expansion in the inflamed intestine. However, this effect might be difficult to point out due to the very low fungal abundance in SPF mouse microbiota. Thus, to go further and reveal a potential phenotype, we induced colitis in mice treated with a broad-spectrum antibiotic and antifongic cocktail and gavaged with *C. albicans* to expand the intestinal fungal load (Fig. 4D). The increased colitis severity in *Card9*^Mrp8cre^ mice was maintained in this setting (Fig. 4E-G). *C. albicans* load was slightly increased (although not statistically significant) in the caecal content (Fig. 4H-I), colon and caecal tissues (Fig. 4J) of *Card9*^Mrp8cre^ mice, and the number of cfu recovered from liver, spleen and kidney were significantly higher compared to *Card9*^Mrp8wt^ mice (Fig. 4K). These results show that *Card9* expression in neutrophils is crucial to control the fungal load in the inflamed gut, the direct translocation of *C. albicans* from the gut to the liver, and to avoid its systemic dissemination.

### The absence of *Card9* impacts neutrophils survival by increasing apoptosis

To explore the mechanisms underlying the impaired capacity of *Card9*^Mrp8cre^ neutrophils to kill *C. albicans* despite intact phagocytosis, autophagy, and ROS production, we investigated neutrophils survival rates by flow cytometry using an AnnexinV-FITC assay coupled to Live/Dead staining (Fig. 5). AnnexinV reveals ongoing apoptosis, whereas Live/Dead only stains permeable dead cells. Interestingly, after 1h incubation at 37°C, *Card9*^Mrp8cre^ neutrophils showed a significant increase in AnnexinV-FITC MFI compared to *Card9*^Mrp8wt^ neutrophils (Fig. 5A-B). Consistently, an increase in percentages of apoptotic (Q1, AnnexinV^+^LD^-^) and dead (late apoptotic/necrotic) neutrophils (Q2, AnnexinV^+^LD^+^), and a decrease in viable neutrophils (Q4, AnnexinV^-^LD^-^), were observed for the *Card9*^Mrp8cre^ genotype (Fig. 5C). Interestingly, the surface expression of the CD62L marker (a surface protein that is lost upon cell activation) was reduced in *Card9*^Mrp8cre^, with a lower percentage of CD62L^+^ cells and a higher percentage of CD62L^-^ cells (Fig. 5D). These results suggest an excessive basal activation of *Card9*^Mrp8cre^ compared to *Card9*^Mrp8wt^ neutrophils. Similar results were obtained with *Card9*^-/-^ versus Card9WT neutrophils (Fig. S5A-D), confirming the impact of *Card9* on neutrophil survival.

**Figure 5.**
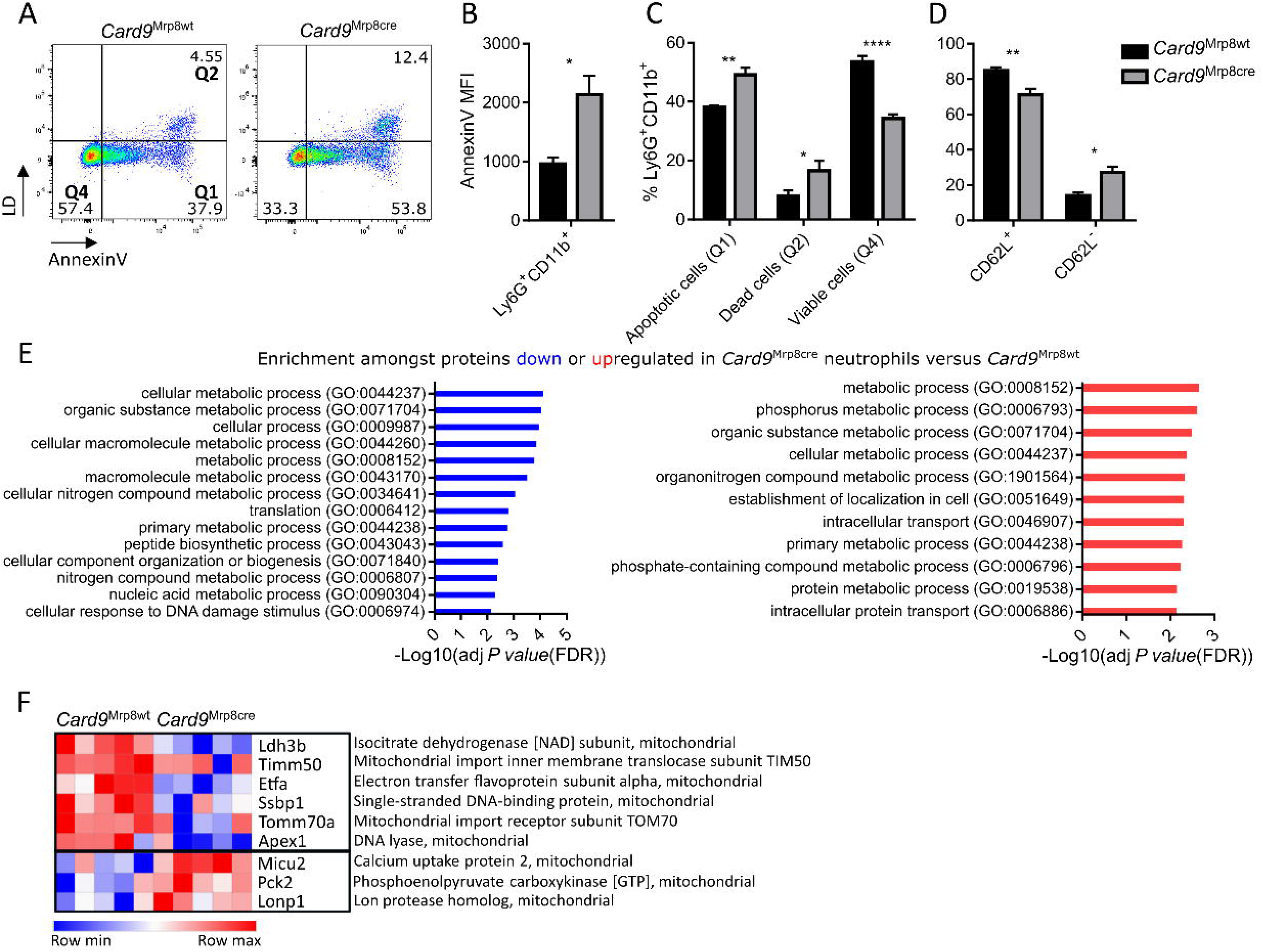
The absence of *Card9* impacts neutrophils survival by increasing apoptosis. (A) Representative flow cytometry plots of Ly6G^+^CD11b^+^ neutrophils purified from bone marrow of *Card9*^Mrp8wt^ (left) and *Card9*^Mrp8cre^ (right) mice, co-stained with AnnexinV and a Live/Dead marker. (B) AnnexinV MFI of Ly6G^+^CD11b^+^ neutrophils from *Card9*^Mrp8wt^ and *Card9*^Mrp8cre^ mice, incubated for 1h at 37°C. (C) Percentage of apoptotic neutrophils (Q1: AnnexinV^+^LD^-^ cells), dead (late apoptotic/necrotic) neutrophils (Q2: AnnexinV^+^LD^+^ cells) and viable neutrophils (Q4: AnnexinV^-^LD^-^ cells) amongst the Ly6G^+^CD11b^+^ population. (D) Percentage of CD62L^+^ and CD62L^-^ neutrophils amongst the Ly6G^+^CD11b^+^ population. Data represent one out of two independent experiments. (E) Histogram representing Gene Ontology biological processes significantly enriched amongst proteins down- (blue) or up-regulated (red) in *Card9*^Mrp8cre^ compared to *Card9*^Mrp8wt^ neutrophils in the unstimulated condition. (F) Morpheus heat-map representing mitochondria-related proteins significantly down- or up-regulated in *Card9*^Mrp8cre^ compared to *Card9*^Mrp8wt^ neutrophils in the unstimulated condition. E and F obtained from proteomics data analysis. *P<0.05, **P<0.01, ***P<0.001, ****P<0.0001 as determined by Mann-Whitney test (B) and two-way ANOVA with Sidak’s post-test (C-D).

To go further, we performed a proteomic analysis on *Card9*^Mrp8cre^ versus *Card9*^Mrp8wt^ neutrophils after 1h incubation at 37°C. This approach revealed that a large number of proteins related to cellular metabolism pathways were differentially regulated between both genotypes. Indeed, a Gene Ontology (GO) functional analysis showed the enrichment in numerous cellular metabolic processes, both in unstimulated and stimulated conditions, among proteins statistically down- or up-regulated in *Card9*^Mrp8cre^ compared to *Card9*^Mrp8wt^ neutrophils (Fig. 5E and S5E). Especially, we observed a high prevalence of mitochondrial proteins among the differentially regulated candidates between the two genotypes (Fig. 5F), suggesting that neutrophil mitochondrial functions are impacted by the absence of *Card9*.

### *Card9* controls neutrophil survival by affecting mitochondrial function

Subsequently, we analyzed neutrophil mitochondrial function using MitoTracker Green (reflecting mitochondrial mass) and MitoTracker Red (reflecting mitochondrial membrane potential) markers in a flow cytometry assay (Fig. 6A). No difference was observed in terms of MFIs, but we observed a significant increase in the percentage of MitoGreen^+^MitoRed^-^ neutrophils (with dysfunctional or metabolically inactive mitochondria) in the *Card9*^Mrp8cre^ compared to the *Card9*^Mrp8wt^ genotype (Fig. 6A). Tetramethylrhodamine methyl ester (TMRM) assay evaluating the mitochondrial membrane potential confirmed the increase in apoptotic, metabolically stressed cells (TMRM^-^) among *Card9*^Mrp8cre^ neutrophils (Fig. 6B). These results suggest that the survival defect of *Card9*^Mrp8cre^ neutrophils may be due to an altered energetic metabolism. Real-time bioenergetic profile analysis using Seahorse technology showed that *Card9*^Mrp8cre^ neutrophils have a higher basal Oxygen Consumption Rate (OCR) during a cell mito stress assay (Fig. 6C), indicating a higher oxidative phosphorylation activity compared to *Card9*^Mrp8wt^ neutrophils. Basal respiration and ATP production rate were both highly increased in *Card9*^Mrp8cre^ neutrophils (Fig. 6D). Moreover, a Seahorse real-time ATP rate assay demonstrated that *Card9*^Mrp8cre^ neutrophils have an increased mitochondrial ATP (mitoATP) production rate, whereas glycolytic ATP (glycoATP) production rate is only mildly decreased, as shown by the ATP rate index and the energetic map (Fig. 6E). A Seahorse glycolytic rate assay confirmed the moderate decrease of the basal Extracellular Acidification Rate (ECAR) of *Card9*^Mrp8cre^ neutrophils (Fig. S6A). However, metabolomics analysis on neutrophil supernatants incubated for 24h revealed a decreased in lactate production, leading to a reduced lactate/glucose ratio in *Card9*^Mrp8cre^ neutrophils (Fig. S6B). Similar results were obtained when we compared Card9WT and *Card9*^-/-^ neutrophils in the assays described above (Fig. S6C-E). Thus, *Card9* deletion in neutrophils induces an overactivation of mitochondria and tends to reduce glycolytic activity, which is the major energy source of normal neutrophils. Blocking glycolysis with 2DG (2-Deoxy-D-glucose) increased the apoptotic rate of *Card9*^Mrp8wt^ but not of *Card9*^Mrp8cre^ neutrophils; showing that glycolysis is more essential to *Card9*^Mrp8wt^ than *Card9*^Mrp8cre^ neutrophils (Fig.6F). Conversely, blocking the mitochondrial respiratory chain with oligomycineA increased the apoptotic rate of *Card9*^Mrp8cre^ but not of *Card9*^Mrp8wt^ neutrophils, demonstrating the crucial role of mitochondria as an energy source in *Card9*^Mrp8cre^ neutrophils (Fig. 6G). Thus, contrary to *Card9*^Mrp8wt^ neutrophils that mainly rely on glycolysis as a source of energy, *Card9*^Mrp8cre^ neutrophils present an altered metabolism with an overactivation of mitochondria associated with a dysfunctional state, leading to apoptosis. Intestinal inflammation is associated with a high degree of oxidative stress. To evaluate the effect of oxidative stress on the phenotype of *Card9*^Mrp8cre^ neutrophils, they were treated for 1h with H_2_O_2_. The increased apoptosis and necrosis rates observed in *Card9*^Mrp8cre^ neutrophils were stronger in oxidative stress than in basal condition (Fig. 6H). These results show that the role of CARD9 in the survival capacity of neutrophils, especially in oxidative conditions, is mediated by effects on mitochondrial functions.

**Figure 6.**
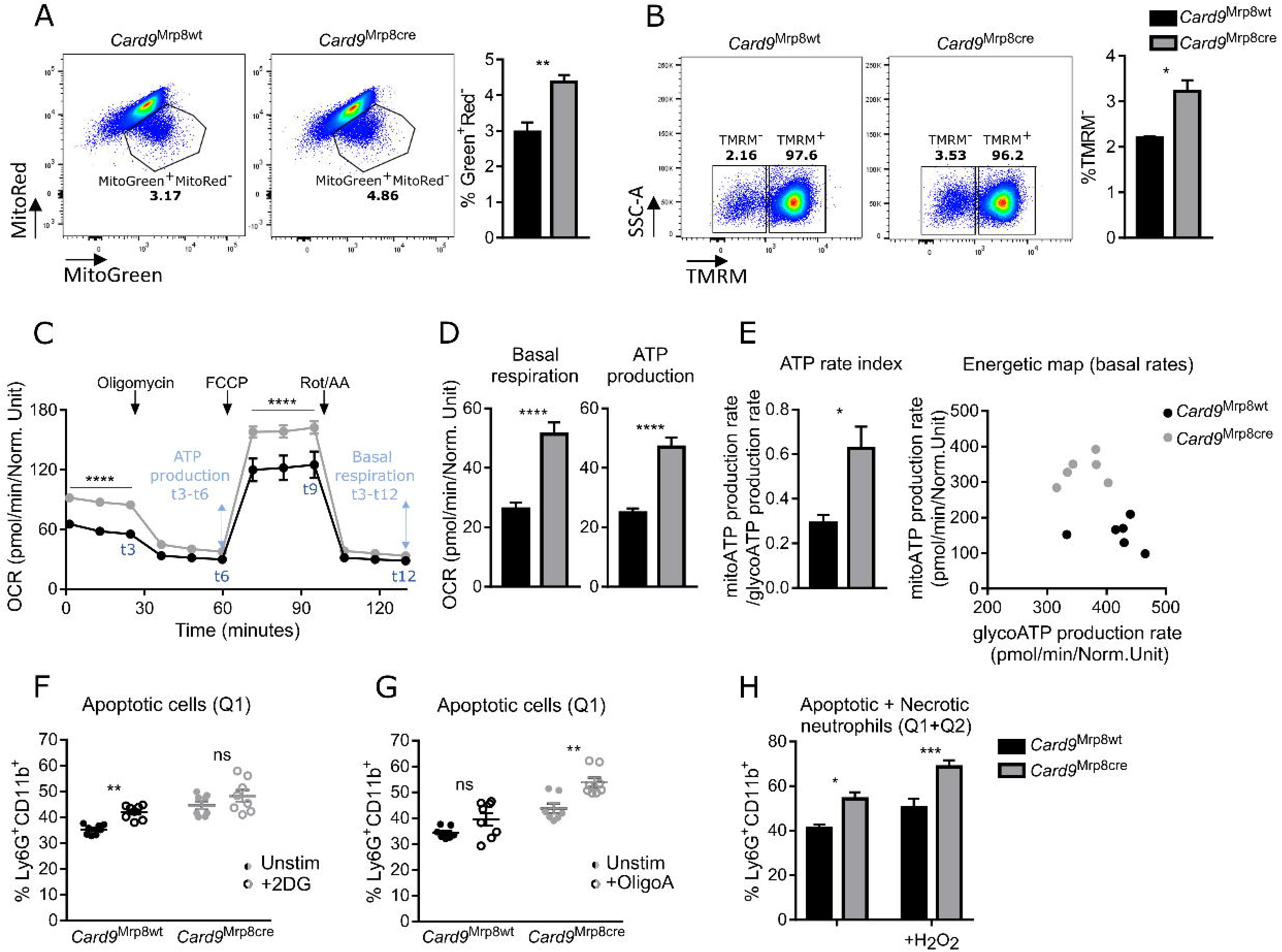
*Card9* controls neutrophil energetic metabolism by affecting mitochondrial function. (A) Representative flow cytometry plots and precentages of MitoGreen^+^MitoRed^-^ cells (corresponding to cells with dysfunctional mitochondria) amongst Ly6G^+^CD11b^+^ neutrophils. Neutrophils were purified from the bone marrow of *Card9*^Mrp8wt^ or *Card9*^Mrp8cre^ mice, and incubated for 1h at 37°C. (B) Representative flow cytometry plots and percentages of TMRM^-^ cells (corresponding to cells with dysfunctional mitochondria/apoptotic or metabolically inactive cells) amongst Ly6G^+^CD11b^+^ neutrophils. (C) Oxygen Consumption Rate (OCR) of *Card9*^Mrp8wt^ and *Card9*^Mrp8cre^ neutrophils measured during a Seahorse Cell Mito Stress assay. (D) Basal respiration (late rate measurement before oligomycin injection (t3) – non-mitochondrial respiration rate (t12)) and ATP production rate (late rate measurement before oligomycin (ATP synthase blocker) injection (t3) - minimum rate measurement after oligomycin injection (t6) obtained from the Seahorse Cell Mito Stress assay. (E) ATP rate index and energetic map of *Card9*^Mrp8wt^ and *Card9*^Mrp8cre^ neutrophils obtained from the Seahorse Real-time ATP rate assay. (F) Percentage of apoptotic cells amongst the Ly6G^+^CD11b^+^ neutrophil population of *Card9*^Mrp8wt^ and *Card9*^Mrp8cre^ genotypes after treatment with 2-DG 10 mM or (G) oligomycin 1.5 μM for 1h. (H) Percentage of apoptotic and necrotic cells amongst the Ly6G^+^CD11b^+^ neutrophil population of *Card9*^Mrp8wt^ and *Card9*^Mrp8cre^ genotypes after addition of H2O2 0.01 mM for 1h to increase oxydative stress. Data are mean ± SEM of at least two independent experiments. *P<0.05, **P<0.01, ***P<0.001, ****P<0.0001 as determined by Mann-Whitney test (A, B, D, E, F) or two-way ANOVA with Sidak’s post-test (G, H). Norm. Unit, normalized unit; FCCP, Carbonyl cyanide 4-(trifluoromethoxy)phenylhydrazone; Rot/AA, Rotenone/AntimycinA; 2DG, 2 deoxy-glucose; OligoA, oligomycinA.

## DISCUSSION

Altogether, this study reveals that, in the absence of CARD9, mitochondrial dysfunction in neutrophils leads to their premature death through apoptosis, especially in an oxidative environment. The decrease of functional neutrophils in the gut affects fungal containment and increases susceptibility to intestinal inflammation. CARD9 polymorphisms are associated with IBD. Mouse studies showed the contribution of CARD9 to host defense and intestinal barrier, notably through the production of IL-22 and the modulation of the gut microbiota metabolic activity^25,26^. However, the role of CARD9 in disease pathogenesis has not been elucidated at the cellular level. Our study reveals that *Card9* deletion in neutrophils, contrarily to epithelial or CD11c^+^ cells, increases the susceptibility to DSS-induced colitis. We thus focused on studying the role of CARD9 expression in neutrophil functionality. Indeed, neutrophils have been studied in different models of IBD and fungal infection, but their direct contribution to pathogenesis and the role of CARD9 in these mechanisms remain poorly understood^31,4,5,32^.

Human CARD9 deficiency results in impaired neutrophil fungal killing, leading to a selective defect to contain invasive fungal infection^33^. In both humans and mice, CARD9 is required in microglia for neutrophil recruitment and control of fungal infection in the central nervous system^34,35^. CARD9 signaling was also involved in neutrophil phagocytosis and NETosis (Neutrophil extracellular traps) functions, enhancing mouse survival to a lethal dose of *C. albicans*^36^. Moreover, CARD9 expression in neutrophils promotes autoantibody-induced arthritis and dermatitis in mice^37^, and inflammation levels in a mouse model of neutrophilic dermatosis^38^. In line with these studies, we found that *Card9* deletion affects the capacity of neutrophils to kill fungi *in vitro* and *in vivo*, with no impact on neutrophil structure, ROS production, autophagy or phagocytosis. We have not examined the role of CARD9 in chemotaxis because, in the context of fungal infection in patients with CARD9 deficiency, neutrophil-intrinsic chemotaxis was not affected^24^. In DSS-induced colitis, *Card9* deficiency in neutrophils does not impact their recruitment to the colon, but reduces the number of mature neutrophils. Indeed, we discovered that the absence of CARD9 increases apoptosis rates in neutrophils, and that this premature death was caused by mitochondria overactivation. Indeed, the intrinsic pathway of apoptosis is initiated by the permeabilization of mitochondria, which releases proapoptotic factors into the cytosol^39,40^. Other CARD proteins mediate apoptotic signaling through CARD-CARD domain interactions^13^. In addition to mediating inflammation, CARD9, a member of the CARD proteins family, was recently shown to inhibit mitochondria-dependent apoptosis of cardiomyocytes under oxidative stress^41^. Here, we show that CARD9 also mediates mitochondrial function and apoptosis in neutrophils, especially in an oxidative environment. Further investigation is required to fully elucidate the impact of mitochondria overactivation on neutrophil function, especially *in vivo*. The impact of *Card9* on neutrophil mitochondrial function might not fully explain the increased susceptibility of *Card9*^Mrp8cre^ mice to DSS colitis. Moreover, other cell types are likely involved in the *Card9*^-/-^ mice phenotype, explaining the impact of *Card9* deletion on microbiota metabolic activity, as we show that this is not dependent on neutrophils^25,26^.

Neutrophils contain very few mitochondria compared to other leukocytes, and depend mainly on glycolysis to produce ATP, which is essential to perform their designated tasks. This allows energy generation in a low-oxygen environment and keeps oxygen available for neutrophil effector functions^42^. Thus, the dependence of neutrophils on glycolysis could be an adaptation to allow oxygen to be used in the anti-microbial response rather than entering oxidative phosphorylation^42^. Here, we show that in the absence of CARD9, mitochondria are overactivated in neutrophils, leading to their premature death and the loss of their anti-microbial functions. Alternatively, excessive oxidative phosphorylation could also lead to an overactivation of neutrophils by increasing ATP production, especially in inflammatory environments where their activity is typically tightly controlled by the low oxygen pressure. Neutrophil overactivation could damage the surrounding tissues and exacerbate inflammation, explaining the increased susceptibility to intestinal colitis in the absence of *Card9*. In humans, the « glycogen storage disease type Ib » induces a functional defect of neutrophils due to glycolysis dysfunction and impaired energy homeostasis^43^. This disease is associated with IBD-like phenotypes, highlighting the importance of neutrophil metabolism in intestinal health^44,43^. Immunometabolism is a central concept as immunity and metabolism impact each other in both ways : (i) energetic metabolism impacts immune function, which is well-documented for lymphocytes and macrophages^45^, but not for neutrophils; and (ii) we show that a protein known for its roles in innate immunity, CARD9, can also impact cell metabolism. Further investigation is required to understand how CARD9 does interact with mitochondrial function in a direct or indirect manner.

Neutrophils are involved in various diseases, including infection, cardiovascular diseases, inflammatory disorders and cancer, which makes them exciting targets for therapeutic intervention^46^. Despite the complex implication of neutrophils in disease, various therapeutic approaches aim to enhance, inhibit or restore neutrophil function, depending on the pathology. In inflammatory diseases with excessive neutrophil activity, their attenuation could be desired, even though killing functions against microorganisms might still need to be preserved. Our work shows that increased apoptosis in neutrophils does not alleviate intestinal inflammation, even though these cells are known to contribute to disease development. Recent studies indicate substantial phenotypic and functional heterogeneity of neutrophils^46^. Thus, targeting a specific subpopulation may allow the attenuation of harmful aspects of neutrophils without compromising host defence. Some neutrophil-targeted therapeutic strategies have reached the clinic, notably in the context of IBD with positive effects of the growth factor granulocyte colony-stimulating factor G-CSF^47^. It opens promising options for numerous complex pathologies.

## Author Contributions

C.D and H.S. conceived and designed the study, performed data analysis, and wrote the manuscript; C.D. designed and conducted all experiments, unless otherwise indicated; C.M., J.P., A.M., A.A., M.-L.M., B. L., G.D.-C., M.S., C. G., M.P., Y.W., A.L. provided technical help for the *in vitro* and/or *in vivo* experiments; J.S. and B.M. performed the immunofluorescence microscopy experiments; C.Pi. and S.C. conducted the proteomics analyses; F.B., E.C. and A.R. performed the metobolomics analysis; T.L. provided mice and C.O. performed genotyping; C.Pe., S.L. and J.E.-B. helped with ROS production assays; R.J.A provided Scenith kits for tests with neutrophils; C.D., C.M., J.P., M.L.-R., P.L., N.R., J.E.-B., B. M. and H.S. discussed experiments and results.

## Acknowledgments

We thank M. Thierry Meylheuc and Ms. Christine Longin from the imaging facility at the microscopy and imaging platform MIMA2 (MIMA2, INRAE, 2018. Microscopy and Imaging Facility for Microbes, Animals and Foods, https://doi.org/10.15454/1.5572348210007727E12, Jouy-en-Josas, France) for precious help in SEM and TEM; the members of the IERP (INRAE) and PHEA (CRSA) animal facilities, and of the @bridge histology plateform (Université Paris-Saclay, INRAE, AgroParisTech, GABI); and Catherine Brenner for the use of Seahorse XF device (IGR). Thanks to BioRender for the graphical abstract. Funding was provided by the ANR-17-CE15-0019-01 grant and a Marie Skłodowska-Curie Actions fellowship.

## Material and Methods

### Mice

*Card9^-/-^, Rag2*^-/-^x*Card9^-/-^, Rag2^-/-^, Card9*loxxMrp8cre, *Card9*loxxVillincre, and *Card9*loxxCD11ccre mice on C57BL/6J background were obtained from the Saint-Antoine Research Center and housed at the IERP (INRAE, Jouy-en-Josas) under specific pathogen-free conditions. Animal experiments were performed according to the institutional guidelines approved by the local ethics committee of the French authorities, the ‘Comité d’Ethique en Experimentation Animale’ (COMETHEA, CEEA45).

### Induction of DSS colitis and colonization with *C. albicans*

Mice were administered drinking water supplemented with 2-3% (wt/vol) DSS (MP Biomedicals) for 5-7 days (depending on colitis severity of each experiment), and then water only for 5d. Animals were monitored daily for weight loss. For *C. albicans* colonization, mice were treated with 0.4mg/ml streptomycin, 300U/ml penicillin G and 0.125mg/ml fluconazole as indicated on Fig. 4.

### Histology

Colon samples were fixed, embedded in paraffin and stained with hematoxylin and eosin. Slides were scanned and analyzed to determine the histological score (Table S1)^25^.

**Table S1.**
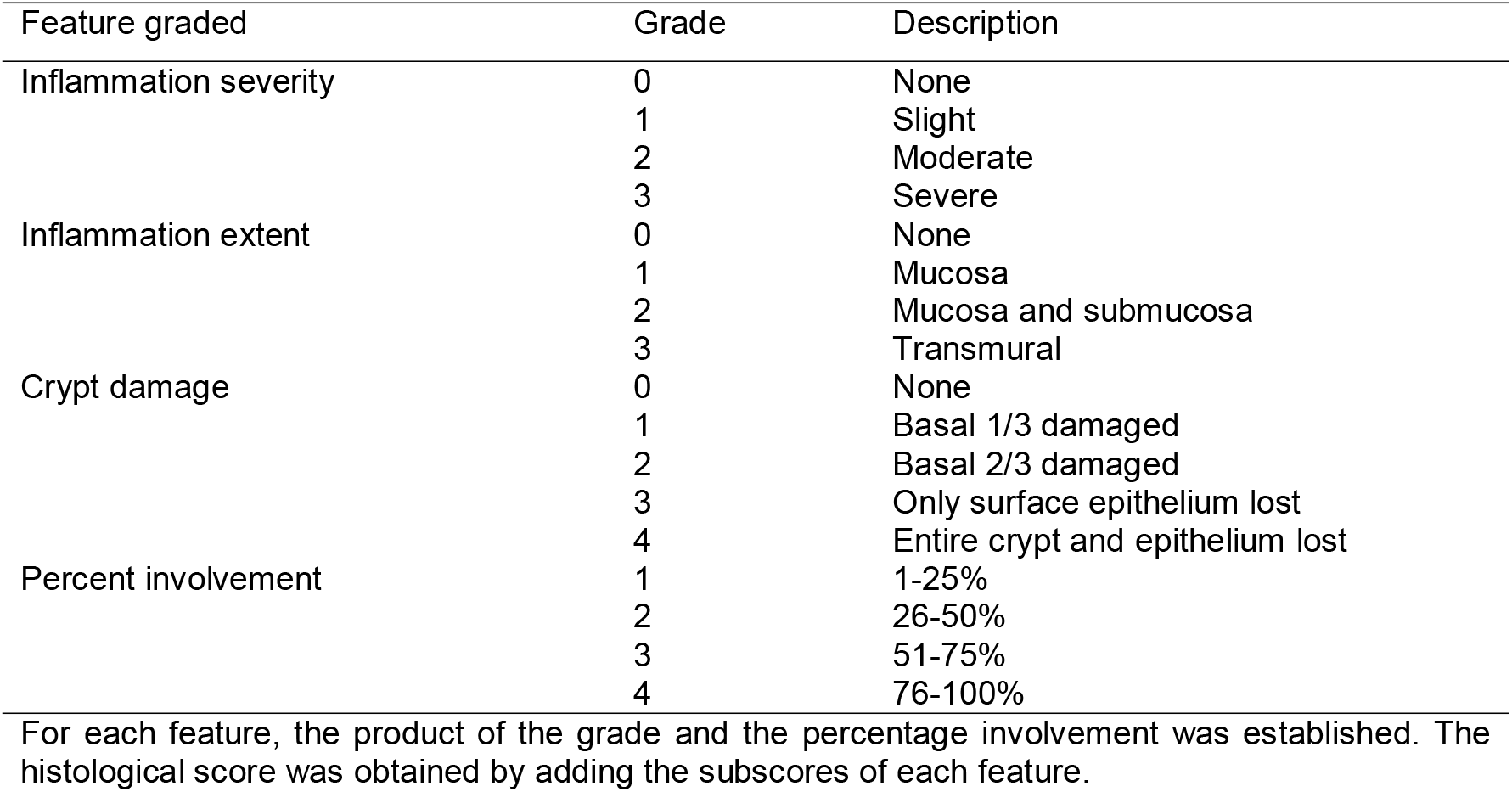
Histological grading of colitis.

### Fecal DNA extraction and total bacteria and fungi quantification

Fecal DNA was extracted as previously described^26^. Luna Universal qPCR Master Mix (New England Biolabs) was used for quantification of fungal ITS2 sequences and TaqMan Gene Expression Assays (Life Technologies) for quantification of bacterial 16S rDNA sequences.

### Cell preparation and stimulation

Epithelial cells were isolated from colonic tissue using a DTT/EDTA buffer. Neutrophils were purified from mouse bone marrow using anti-Ly6G MicroBeads UltraPure and MACS separation columns (Miltenyi Biotec). CD11c^+^ cells were purified from mouse spleen using anti-CD11c MicroBeads UltraPure and MACS separation columns (Miltenyi Biotec). Purity checks and cell counts were performed using a BD Accuri C6 flow cytometer (BD Biosciences). After purification, neutrophils were seeded in 96-well suspension plate (Sarstedt), rested for 30min in RPMI 1640 Medium (Gibco, ThermoFisher Scientific) before addition of 2% heat-inactivated Fetal Bovine Serum (FBS), 2-DG 10mM, oligomycine 1.5μM or H2O2 0.01M and incubated at 37°C as indicated. Isolation of lamina propria immune cells was performed as previously described^48^.

### Gene expression analysis using quantitative RT-PCR

Total RNA was isolated from colon samples or cell suspensions using RNeasy Mini Kit (Qiagen), and quantitative RT-PCR performed using QuantiTect Reverse Transcription Kit (Qiagen) and Luna® Universal RT-PCR Kit (New England Biolabs) in a StepOnePlus apparatus (Applied Biosystems) with specific mouse oligonucleotides (Table S2). We used the 2-ΔΔ^*Ct*^ quantification method with mouse *Gapdh* as an endogenous control and the WT group as a calibrator.

**Table S2.**
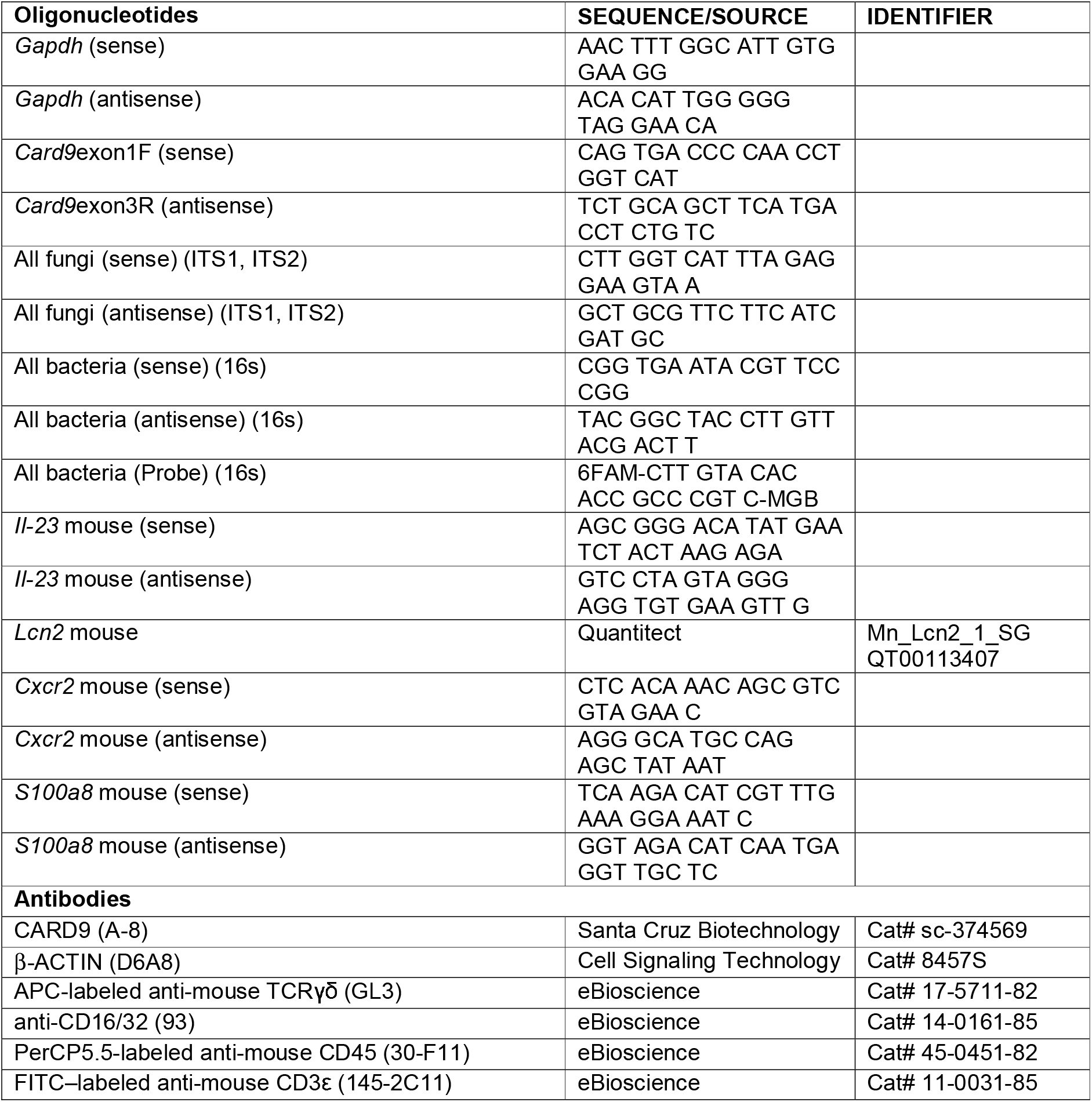
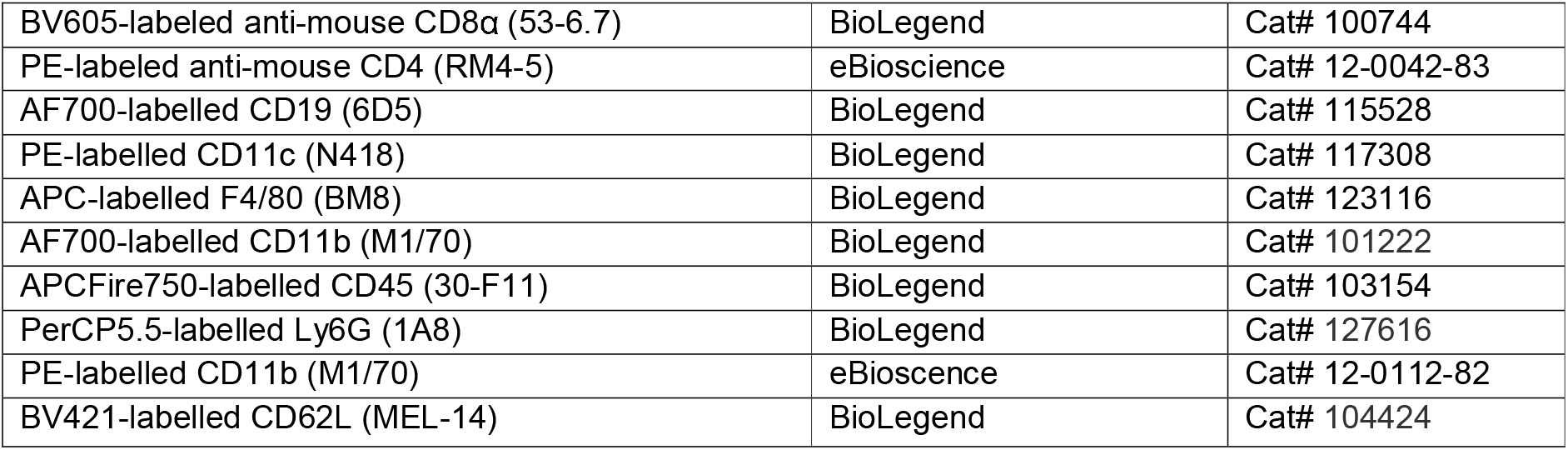
List of oligonucleotides and antibodies.

### Immunoblot Analysis

Mouse tissue or cell suspensions were lysed using Laemmli buffer, loaded on a SDS-PAGE and analyzed with antibodies against CARD9 (A-8:sc-374569, Santa Cruz Biotechnology), or β-ACTIN (D6A8, CST).

### Flow cytometry, cell sorting and functional assays

Flow cytometry was carried out by using LSR Fortessa X-20 (BD) and cell sorting FACS Aria machines (BD). For apoptosis assay, 4×10^5^ neutrophils were stained with AnnexinV-FITC in Binding Buffer (Miltenyi Biotec). For mitochondria analyses, MitoTracker™ Green and MitoTracker™ Red FM were added to neutrophils in MACS buffer for 15min at RT (ThermoFisher Scientific). Alternatively, neutrophils were incubated with TMRM in RPMI for 20min at 37°C (Abcam). For phagocytosis assay, 10^5^ neutrophils stimulated with zymosan-FITC (50μg/ml, Fluorescein zymosanA BioParticles conjugates, FisherScientific), *C. albicans*-GFP (MOI1:1) or *Escherichia coli*-GFP (MOI1:10) for 45min. Cells were stained with surface antibodies in MACS buffer (Table S2).

### Endpoint dilution survival assay

Isolated mouse neutrophils were seeded at 10^5^ cells/well in 96-well plates in RPMI+2% FBS and infected with *C. albicans* (serial fourfold dilutions of an OD=1 solution were added by raws (row1, MOI20:1; row2, MOI5:1…). After 24h at 37°C, colonies were visualized with a Nikon TMS inverted microscope and counted at the lowest dilutions (row7-8).

### Killing assay

Purified neutrophils were seeded in 96-well plate at 10^6^ cells/well and stimulated with PBS, *C. albicans* (MOI1) or *E. coli* (MOI10) for 90min at 37°C. Cells were washed and lysed with 200μl TritonX100 0.025%. Serial dilutions were plated on YEPD or LB plates.

### Oxydative burst

Experiments were performed on a TriStar LB942 Reader using 10^5^ neutrophils in 200μL HBSS (ThermoFisher Scientific) and luminol 80μM (Sigma) and stimulated with phorbol 12-myristate 13-acetate (PMA; 0.1μg/mL; Sigma) or opsonized zymosan (20mg/mL zymosan A from *Saccharomyces cerevisiae*; Sigma). The indexed maximal relative luminescence (in relative light units [RLU]) was calculated as follow: indexed RLU max=([hemochromatosis patient maximal RLU]/[healthy donor maximal RLU])×100. Alternatively, superoxyde dismutase (5Units/mL) and catalase A (10Units/mL) were added to differentiate total and intracellular ROS production. Absorbance was measured at 550nm for 30min.

### Real time bioenergetic profile analysis

Mito Stress Test, Glycolytic Rate Assay and Real-time ATP assay were performed on a XF96 Extracellular Flux Analyzer (Seahorse Biosciences). Mouse neutrophils were seeded at 2×10^5^ cells/well in RPMI+2%FBS in 0.01% poly-L-lysine pre-coated plate. After 2h rest, cells were washed in Seahorse RPMI medium and incubated for 1h at 37°C without CO2. In the analyzer, oligomycin 1.5μM, FCCP 1μM, Rotenone+AntimycinA 0.5μM and 2DG 50μM were injected at the indicated times. Protein standardization was performed after each experiment, with no noticeable differences in protein concentration and cell phenotype.

### Electron microscopy

Purified neutrophils were stimulated with *C. albicans* (MOI 2) or *E. coli* (MOI 10) for 1h at 37°C, washed in PBS and fixed. Samples preparation and SEM/TEM analyses were performed at the Microscopy and Imaging Platform MIMA2 (Université Paris-Saclay, INRAE, AgroParisTech, Jouy-en-Josas, France, https://doi.org/10.15454/1.5572348210007727E12).

### Proteomics

5μg protein extracts were submitted to in-gel digestion. Desalting was performed as described before^49^. Peptides were analyzed on a nanoElute-timsTOF ProLC-MS/MS system (Bruker)^50^. Raw files were analyzed using MaxQuant v1.6.10.43: database UP000000589_10090 (21994 entries, 17-Jun-2020). Data filtering, imputation and statistical analysis were performed with ProStar Zero 1.20.0^50^ Proteins with FDR<5% (Pounds method) were significant with a fold change >1.2.

### Nuclear Magnetic Resonance

200μl culture media were analyzed by 1D 1H-NMR. All NMR spectra were recorded on a Bruker AvanceIII 800MHz spectrometer equipped with a QPCI 5mm cryogenic probe head. Spectra were acquired and processed using the Bruker Topspin 4.0 software. Quantification of glucose and lactate was performed using addition of 25% TSPd4 in D2O as internal standard.

### Statistical analysis

Statistical analysis was performed using GraphPad Prism 7 software (see figure legends).

## Supplemental Figure and Table

**Figure S1. *Card9* is mainly expressed by myeloid cells.** (A) *Card9* expression in different organs of *Card9*WT and *Card9*^-/-^ mice by quantitative RT-PCR analyses, normalized to *Gapdh*. (B) CARD9 and β-ACTIN expression in different organs of *Card9*WT and *Card9*^-/-^ mice by Western Blot analyses. (C) *Card9* expression in different sorted cell populations from the spleen and bone marrow of *Card9*WT and *Card9*^-/-^ mice by quantitative RT-PCR analyses, normalized to *Gapdh*.

**Figure S2. Validation of conditional knockout mice strains.** (A) *Card9* expression in colonic epithelial cells and tissue from *Card9*^Villinwt^ and *Card9*^Villincre^ mice (up right panel), CD11c^+^ purified cells and CD11c fraction from the spleen of *Card9*^Cd11cwt^ and *Card9*^Cd11ccre^ mice (up left panel) and Ly6G^+^ purified neutrophils and Ly6G^-^ fraction from the bone marrow of *Card9*WT, *Card9^-/-^, Card9*^Mrp8wt^ and *Card9*^Mrp8cre^ mice (lower panels) by qRT-PCR analyses, normalized to *Gapdh*. (B) CARD9 and β-ACTIN expression in Ly6G^+^ purified neutrophils and Ly6G^-^ fraction from the bone marrow of *Card9*WT, *Card9^-/-^, Card9*^Mrp8wt^ and *Card9*^Mrp8cre^ mice by Western blot. (C) Flow cytometry plots representing total cell populations of the bone marrow of a C57BL/6 mouse (left panel), and Ly6G^+^CD11b^+^ neutrophils purified from the bone marrow using Ly6G ultrapure magnetic beads and MACS column (right panel). (D) Colon length of DSS-exposed *Card9*^Villinwt^*, Card9*^Villincre^*, Card9*^Cd11cwt^ and *Card9*^Cd11ccre^ mice. (E) AhR activity of feces from *Card9*^Mrp8wt^ and *Card9*^Mrp8cre^ mice at steady-state, using HepG2 reporting cells. (F) Weight and DAI score of DSS-exposed *Card9*^Mrp8wt^ or *Card9*^Mrp8cre^ mice for 9 days.

**Figure S3. Imaging of *Card9*-deleted neutrophils.** (A) Immunofluorescence staining of *Card9*WT, *Card9^-/-^, Card9*^Mrp8wt^ and *Card9*^Mrp8cre^ neutrophils unstimulated or stimulated with zymosan for 1 or 3h. Blue for DNA, red for β-ACTIN. Scale bars, 5 μm. (B) Scanning electron microscopy (SEM, left) or Transmission electron microscopy (TEM, right) of *Card9*^Mrp8wt^ and *Card9*^Mrp8cre^ neutrophils unstimulated or stimulated with *C. albicans* or *E. coli* for 1h. White arrows point *C. albicans* or *E. coli* phagocytosed by neutrophils. Scale bars, 1 or 1.5 μm.

**Figure S4. *C. albicans* killing capacity is impacted by *Card9* deletion in neutrophils, but not phagocytosis capacity, ROS production or autophagy.** (A) Representative flow cytometry plots showing percentages of Ly6G^+^FITC^+^ and Ly6G^+^GFP^+^ *Card9*WT or *Card9*^-/-^ neutrophils stimulated by zymosan-FITC (50μg/ml, Fluorescein zymosan A BioParticles conjugates), *C. albicans-GFP* (MOI 1:1) or *E. coli-GFP* (MOI 1:10) for 45 min. (B) Graph representing phagocytosis rates obtained by flow cytometry analysis. (C-D) Oxydative burst kinetics of *Card9*WT, *Card9^-/-^, Card9*^Mrp8wt^ and *Card9*^Mrp8cre^ neutrophils as measured by luminol-amplified chemiluminescence of *Card9*WT and *Card9*^-/-^ (left panel) and *Card9*^Mrp8wt^ and *Card9*^Mrp8cre^ (right panel) neutrophils after stimulation with (C) PMA (phorbol 12-myristate 13-acetate; 0.1 μg/mL; Sigma) or (D) opsonized zymosan (20 mg/mL, zymosan A from *Saccharomyces cerevisiae* opsonized with 10% SVF). (E) Total and Intracellular Reactive Oxygen Species (ROS) produced by *Card9*WT and *Card9*^-/-^ neutrophils (left), or *Card9*^Mrp8wt^ and *Card9*^Mrp8cre^ neutrophils (right) in unstimulated condition or after 90 min stimulation with non-opsonized zymosan (0.5 mg/mL). RLU, relative light unit. (F) Western blot showing p62 (62 KDa) and LC3BI/LC3BII (16 and 14 KDa) autophagy proteins in *Card9*WT, *Card9^-/-^, Card9*^Mrp8wt^ and *Card9*^Mrp8cre^ neutrophils purified from the bone marrow and incubated 1h in RPMI +2% FBS at 37°C. (G) Fungi/bacteria loads in feces from *Card9*^Mrp8wt^ or *Card9*^Mrp8cre^ mice during DSS colitis measured by qRT-PCR (ratio of 2^-Ct^).

**Figure S5. The absence of *Card9* impacts neutrophils survival by increasing apoptosis.** (A) Representative flow cytometry plots of Ly6G^+^CD11b^+^ neutrophils purified from bone marrow of *Card9*WT (left) and *Card9*^-/-^ (right) mice, co-stained with AnnexinV and a Live/Dead marker. (B) AnnexinV MFI of Ly6G^+^CD11b^+^ neutrophils from *Card9*WT and *Card9*^-/-^ mice, incubated for 1h at 37°C. (C) Percentage of apoptotic neutrophils (Q1: AnnexinV^+^LD^-^ cells), dead (late apoptotic/necrotic) neutrophils (Q2: AnnexinV^+^LD^+^ cells) and viable neutrophils (Q4: AnnexinV^-^LD^-^ cells) amongst the Ly6G^+^CD11b^+^ population. (D) Percentage of CD62L^+^ and CD62L^-^ neutrophils amongst the Ly6G^+^CD11b^+^ population. Data represent one out of two independent experiments. *P<0.05, **P<0.01, ***P<0.001, ****P<0.0001 as determined by Mann-Whitney test (B) or two-way ANOVA with Sidak’s post-test (C-D).

**Figure S6. *Card9* controls neutrophil energetic metabolism by affecting mitochondrial function.** (A) Extracellular Acidification Rate (ECAR) of *Card9*^Mrp8wt^ and *Card9*^Mrp8cre^ neutrophils measured during a Seahorse Glycolytic rate assay in unstimulated condition (RPMI + 2% FBS). Graph represents basal glycolysis (t2). (B) Lactate/glucose ratio in 24h culture supernatant of *Card9*^Mrp8wt^ and *Card9*^Mrp8cre^ neutrophils measured by metabolomics data analyses. (C) Representative flow cytometry plots and percentages of MitoGreen^+^MitoRed^-^ cells (corresponding to cells with dysfunctional mitochondria) amongst the Ly6G^+^CD11b^+^ neutrophil population from *Card9*WT or *Card9*^-/-^ mice incubated for 1h at 37°C. (D) Representative flow cytometry plots and percentages of TMRM^-^ cells (corresponding to cells with dysfunctional mitochondria/apoptotic or metabolically inactive cells) amongst the Ly6G^+^CD11b^+^ neutrophil population incubated for 3h at 37°C. (E) Oxygen Consumption Rate (OCR) and Extracellular Acidification Rate (ECAR) of *Card9*WT or *Card9*^-/-^ neutrophils measured during a Seahorse Cell Mito Stress assay or a Seahorse Glycolytic rate assay, respectively. Data are mean ± SEM of at least two independent experiments. *P<0.05, **P<0.01, ***P<0.001, ****P<0.0001 as determined by Mann-Whitney tests. FCCP, Carbonyl cyanide 4-(trifluoromethoxy)phenylhydrazone; Rot/AA, Rotenone/AntimycinA; 2-DG, 2-deoxy-D-glucose.

