## Supplementary figures and images for "CARD9 in Neutrophils Protects from Colitis and Controls Mitochondrial Metabolism and Cell Survival"

### Supplemental Figure 1

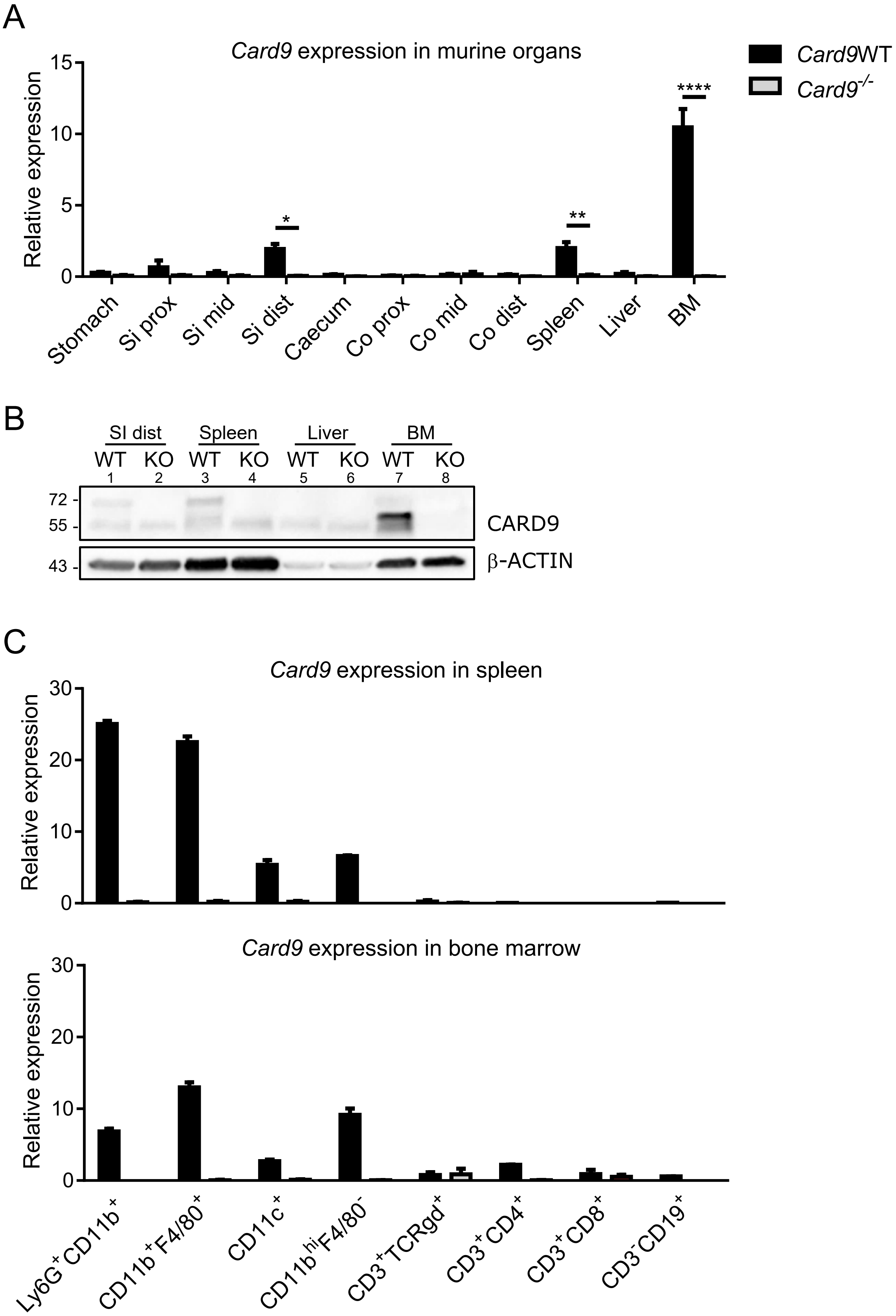

### Supplemental Figure 2

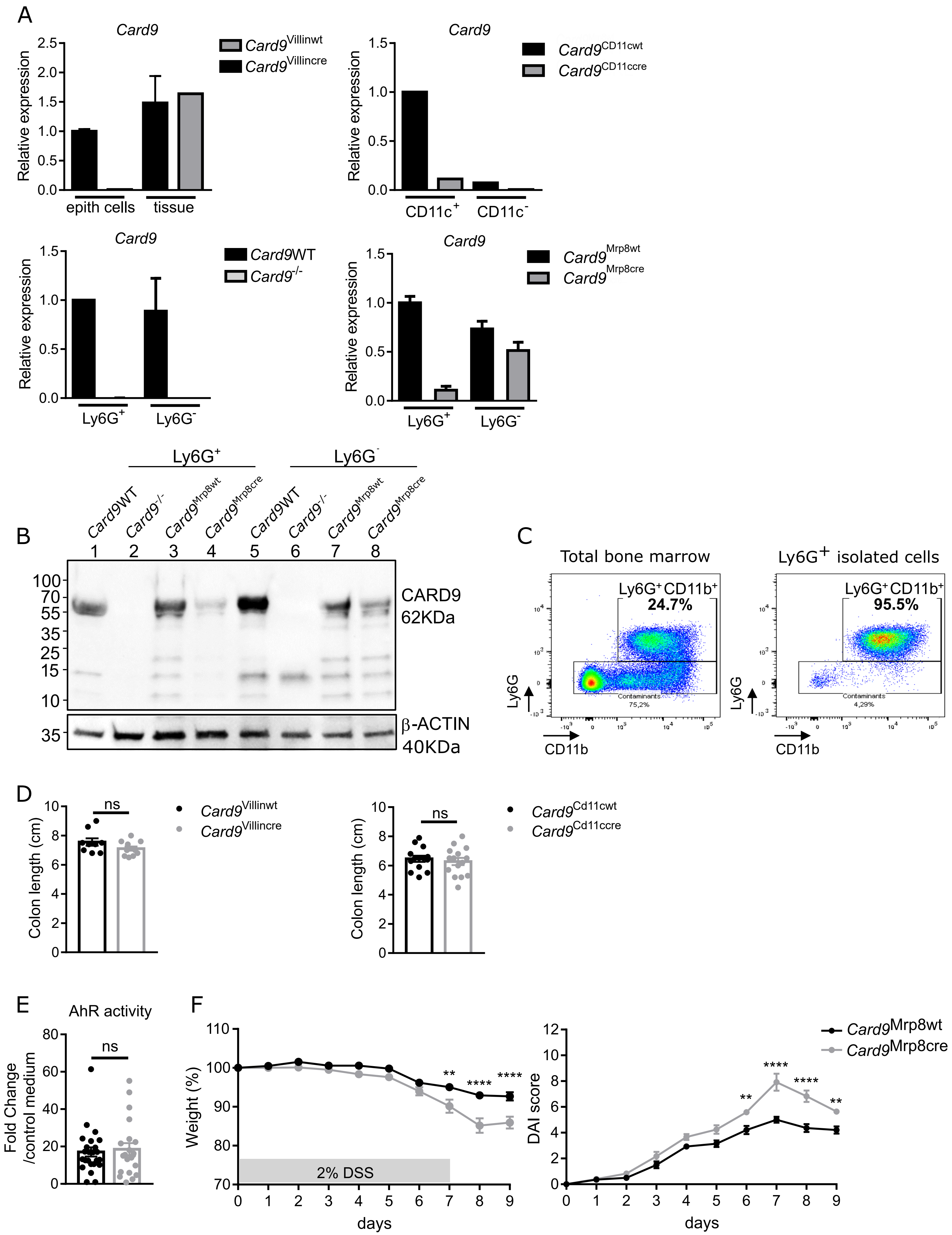

### Supplemental Figure 3

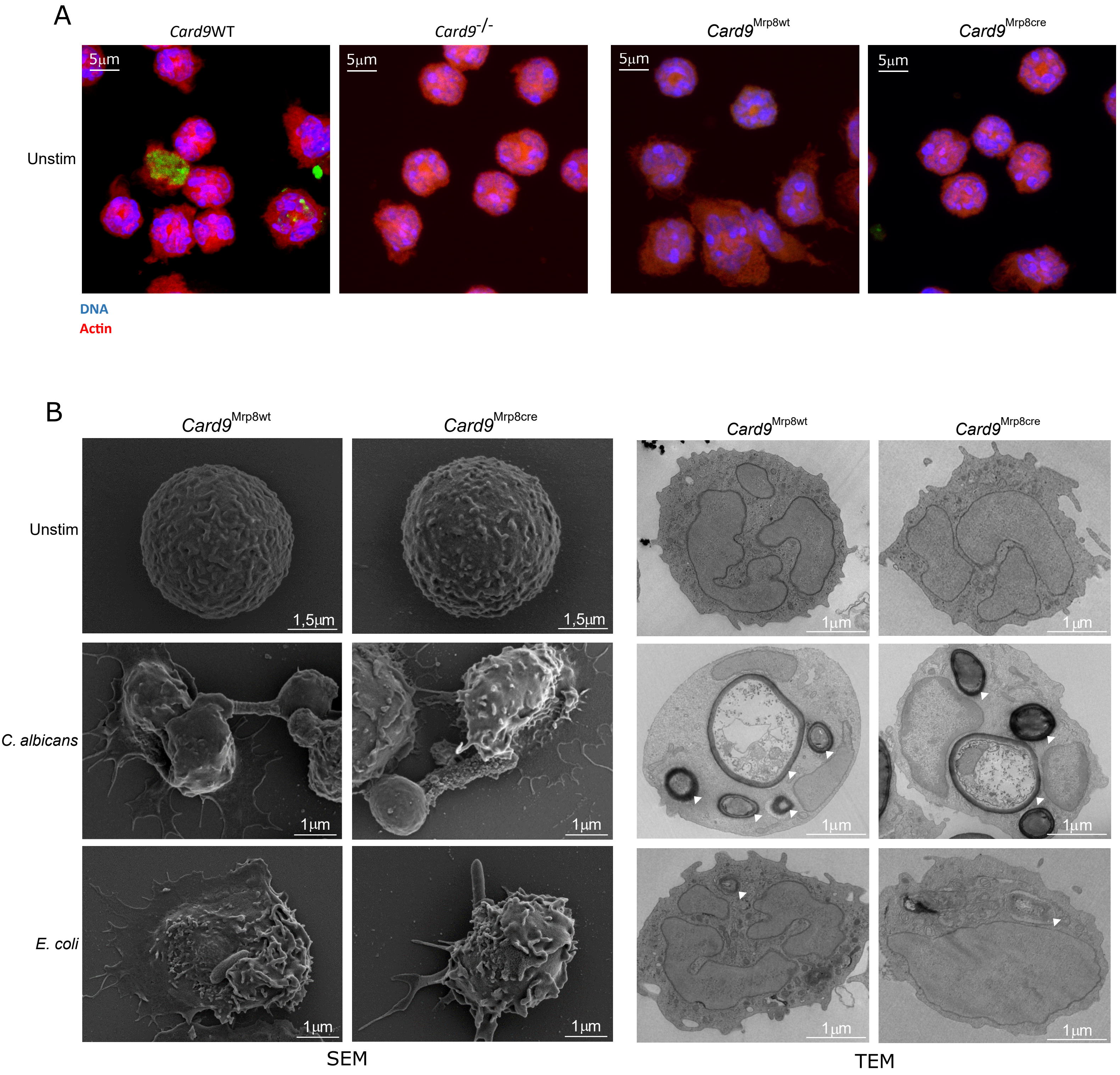

### Supplemental Figure 4

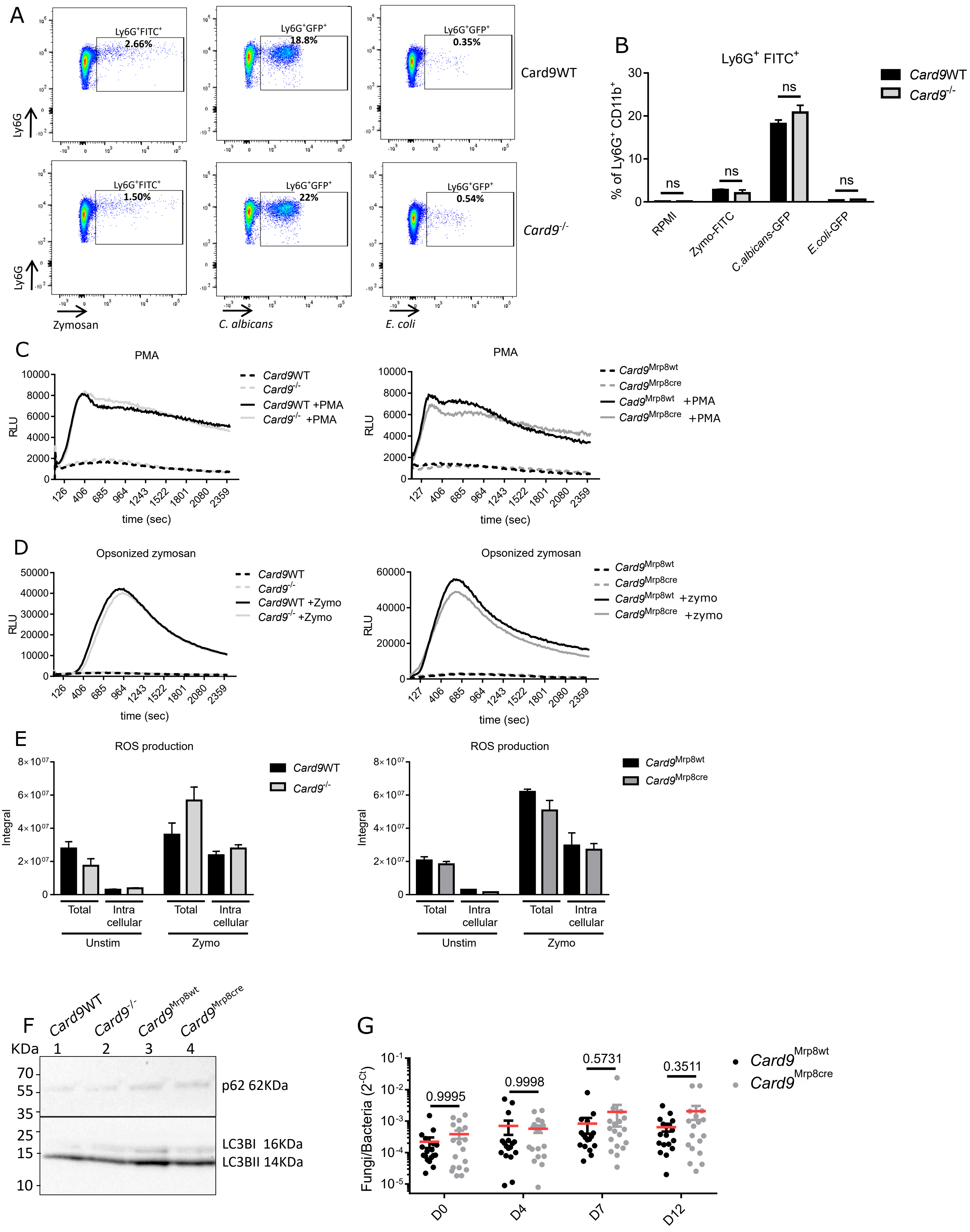

### Supplemental Figure 5

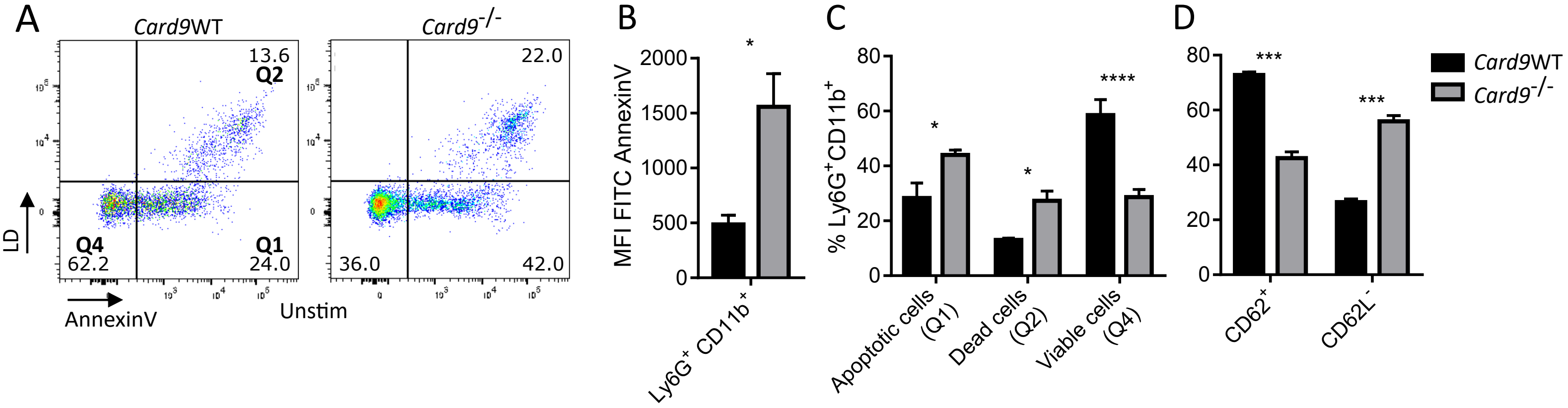

### Supplemental Figure 6

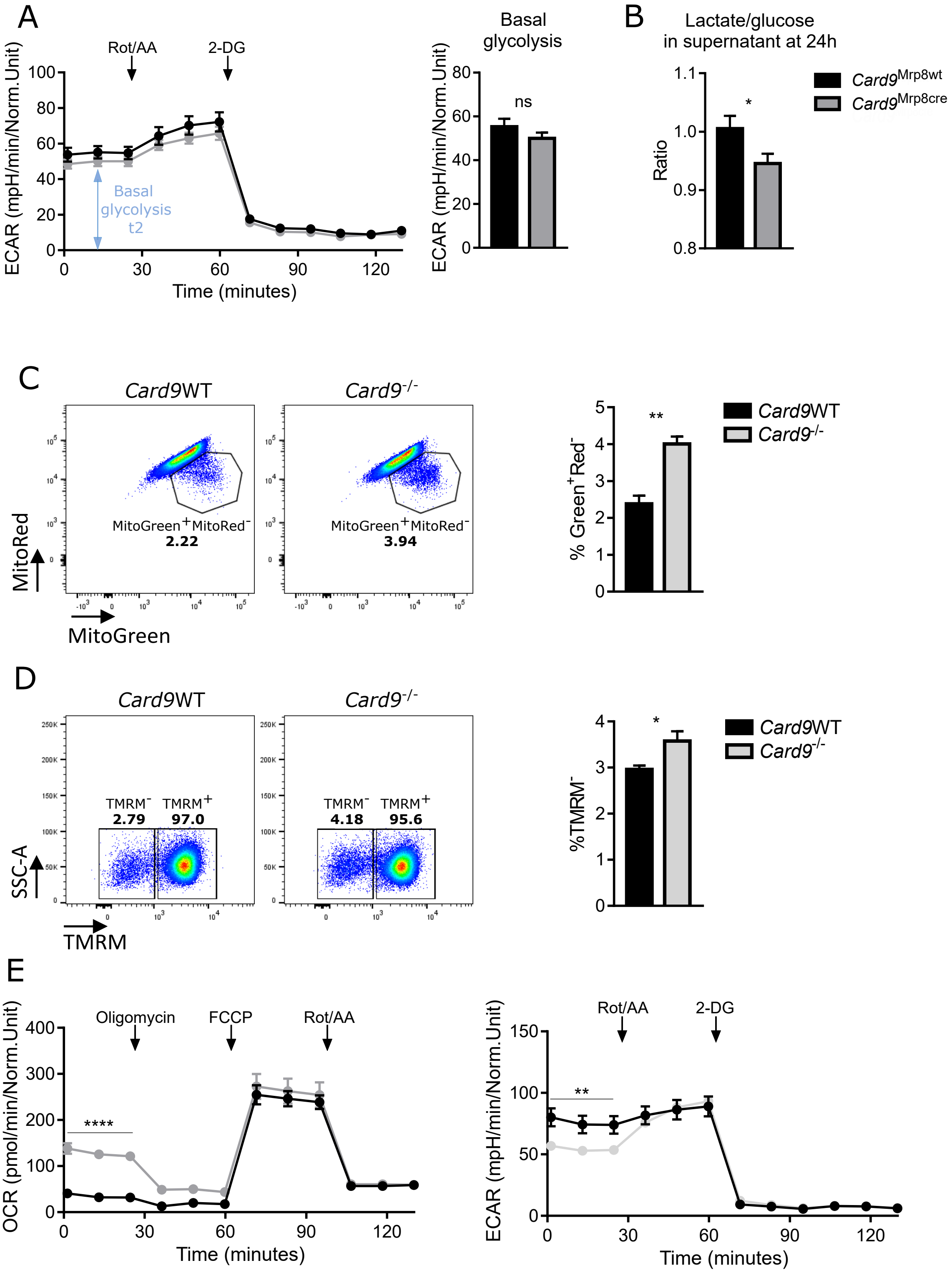
